# ZEB1 shapes AML immunological niches suppressing CD8 T-cell activity while fostering Th17 cell expansion

**DOI:** 10.1101/2023.07.08.548195

**Authors:** Barbara Bassani, Giorgia Simonetti, Valeria Cancila, Marilena Ciciarello, Annamaria Piva, Claudia Chiodoni, Daniele Lecis, Alessandro Gulino, Eugenio Fonzi, Laura Botti, Paola Portararo, Massimo Costanza, Juerg Schwaller, Alexandar Tzankov, Maurilio Ponzoni, Fabio Ciceri, Niccolò Bolli, Antonio Curti, Claudio Tripodo, Mario P. Colombo, Sabina Sangaletti

**Affiliations:** Department of Research, Fondazione IRCCS Istituto Nazionale Tumori, Milan, Italy; IRCCS Istituto Romagnolo per lo Studio dei Tumori (IRST) “Dino Amadori” Meldola, Italy; University of Palermo School of Medicine, Palermo, Italy; Department of Experimental, Diagnostic and Specialty Medicine – DIMES, Institute of Hematology “ Seràgnoli”, Bologna, Italy; Unit of Biostatistics and Clinical Trials, IRCCS Istituto Romagnolo per lo Studio dei Tumori (IRST) “Dino Amadori”, Meldola (FC), Italy; Molecular Neuro-Oncology Unit, Department of Clinical Neuroscience, Fondazione IRCCS Istituto Neurologico Carlo Besta, Milan, Italy; University of Basel Children’s Hospital; Institute of Medical Genetics and Pathology, University Hospital Basel, University of Basel, Basel, Switzerland; IRCCS Ospedale S. Raffaele; University Vita-Salute San Raffaele; Milan, Italy; Hematology Division, Foundation IRCCS Ca’ Granda Ospedale Maggiore Policlinico, 20122 Milan, Italy; Department of Oncology and Hemato-Oncology, University of Milan, 20122 Milan, Italy

**Keywords:** ZEB1, Acute myeloid Leukemia, IL-17, Microenvironment, Bone marrow

## Abstract

Acute myeloid leukemia (AML) development and progression is favored by immune suppression directly triggered by leukemia cells. ZEB1 is a key transcription factor in epithelial-to-mesenchymal transition which, we show here, is capable immune regulation in AML. Leukemic cells which had ZEB1 knocked down have reduced engraftment and extramedullary disease when transplanted into immune competent mice due to concomitant activation of CD8 T lymphocytes and reduced expansion of Th17 cells. Differently, in ZEB1 competent AML, IL-17 sustains the development of a pro-invasive and self-maintaining loop inducing *MMPs* and *SOCS2.* In humans, AML patients show, *in situ* on bone marrow biopsies, a direct correlation between ZEB1 and Th17 and, in gene expression profile when divided according to the median value of *ZEB1* expression, a different overall survival and relapse along with the expression of MMPs, SOCS2 and Th17 cells enrichment. Overall, our data shed new light into the role of ZEB1 in AML that entwines both pro-tumoral and immune regulatory capacity in AML blasts.

## INTRODUCTION

The bone marrow is a peculiar primary lymphoid organ in which the recirculation of naïve T-cell and the presence of antigen-presenting cells competent for presentation may trigger anti-leukemia T-cell responses. On this line, the number of T cells, present in the BM at diagnosis, correlates with overall survival in newly diagnosed acute myeloid leukemia (AML) patients(Lamble et al., 2020) (Davidson-Moncada et al., 2018) (Vadakekolathu et al., 2020). Progression seen in AML suggests that immune suppressive mechanisms should be in place overcoming anti-tumor T-cell response. In the hematopoietic niche, leukemic cells interact with BM stromal cells establishing favorable conditions for survival, proliferation and resistance to therapy as well as escape from immune recognition (Tettamanti et al., 2021) (Mendez-Ferrer et al., 2020). As reported in solid tumors, potentially immunogenic leukemia cells seem to develop multiple mechanisms for immune escape including the establishment of immune suppression. These different immunomodulatory mechanisms encompass regulatory T cells, myeloid-derived suppressor cells (MDSCs) (Han et al., 2014)engagement of inhibitory T-cells pathways (i.e. PD-L1-/PD-1, arginase(Arg)-2 (Li et al., 2020) (Mussai et al., 2019)or interference with specific metabolic pathways through indoleamine-2,3-dioxygenase (IDO (Curti et al., 2007). The engagement of bystander cells, such as BM-MSCs, or a direct endogenous activity of AML blast seem necessary to produce key factors capable of regulating immune cells activities. ZEB1 has been extensively studied in solid cancers as a main transcription factor involved in epithelial-to-mesenchymal transition (EMT) (Caramel et al., 2018).Recent evidences indicate that ZEB1 could regulate also immune cell functions. Indeed, reciprocally to ZEB2, another member of the ZEB family, ZEB1 is expressed by a variety of immune cells (Scott and Omilusik, 2019)with immune suppressive functions. In tumor-associated macrophages ZEB1seems to promote their polarization toward a pro-tumor phenotype (Cortes et al., 2017), also its acts as repressor of miR-200, which negatively regulates the expression of the PD-L1 immune checkpoint (Chen et al., 2014). As mutated counterparts of normal myeloid cells (Mussai *et al*., 2019). AML blasts could adopt ZEB1 expression to modulate the leukemia microenvironment. The expression of ZEB1 has been already reported in leukemia leading to different and, sometimes, opposite conclusions, being ZEB1 described as both pro- and anti-leukemogenic (Shousha et al., 2019) (Stavropoulou et al., 2016) (Li et al., 2021).

In this study, we functionally explored the impact of ZEB1 in murine leukemia cells seeding the BM microenvironment and confirmed data and clinical relevance in AML patients.

## MATERIALS AND METHODS

### Stable gene-silencing

Lentiviral Particles were purchased from OriGene Technologies (catalog number TL513177V). Two specific constructs (“seq-C” and “seq-D”) were tested for efficiency compared to a negative control construct (“Scr”). For ZEB1 stable silencing in human K562 cells, we used the Mission Lentivirus Transduction Particles (pLKO.1-shZEB1-565 - TRCN0000017565 and pLKO.1-shZEB1-631 - TRCN0000364631), purchased from Sigma-Aldrich. A non-target (Scramble - SHC003) sequence was used as negative control.

### Total RNA extraction, reverse transcription, and quantitative polymerase chain reaction (qPCR)

Total RNA was extracted using the Quick RNA micro prep kit (Zymo Research) and subsequently quantified by NanoDrop 2000c Spectrophotometer (Thermo Scientific). cDNA was generated using the high capacity Reverse Transcriptase kit (Applied Biosystems) according to the manufacturer’s instructions. Taqman Probes used are listed in **Supplementary Table 1**. Values were normalized to internal control (β-Actin) using the ΔCT method. For IL-17A stimulation experiments, 5×10^4^ human and murine cell lines expressing or silenced were treated for 7 days with 50ng/mL rIL-17A. Untreated cells were used as control. ΔΔCT results are shown. rIL-17A-stimulated cells were normalized to untreated cells.

#### Gene expression profiling on murine cells

Gene expression profiles were established by Thermo Fisher Mouse Clariom S Assay. RNA labeling, processing, and hybridization were performed according to manufacturer’s instructions, and microarrays were scanned with the Gene Chip System 3000 scanner. Raw data were pre-processed using the sst-RMA algorithm implemented in the Transcriptome Analysis Console software (Thermo Fisher). Downstream analyses were performed on pre-processed data using R software. Multiple probes representing the same gene were collapsed by selecting the probe with the highest variance across samples through the collapseRows function in the WGCNA package (Miller et al., 2011). Differentially expressed genes (DEG) were identified using limma package (Phipson et al., 2016). P-values were adjusted for multiple testing using the Benjamini–Hocheberg false discovery rate (FDR). Genes with an FDR<0.05 were considered statistically significant. Gene set enrichment analysis (GSEA (Subramanian et al., 2005)) was carried out in pre-ranked mode, according to the limma t-statistic, using the hallmark gene set collection from the MSigDB database (http://software.broadinstitute.org/gsea/msigdb/index.jsp). Gene sets with an FDR<0.05 were considered statistically significant.

#### Human gene expression datasets

Public gene expression data were obtained from the Gene Expression Omnibus (GEO), repository (https://www.ncbi.nlm.nih.gov/gds): GSE6891 (Verhaak et al., 2009), GSE12417, GSE15434, GSE16015, GSE37642 (Li et al., 2013), GSE66525 (Hackl et al., 2015). NGS-PTL (Next Generation Sequencing platform for targeted Personalized Therapy of Leukemia) array data from 61 AML bone marrow samples (blasts ≥80%) have been previously generated (Simonetti et al., 2021) (GSE161532) and patients characteristics are listed in **Supplementary Table 2**. The Beat AML (Tyner et al., 2018)and The Cancer Genome Atlas project on AML (Cancer Genome Atlas Research et al., 2013) transcriptomic cohorts were obtained from https://portal.gdc.cancer.gov/ (projects BEATAML1.0-COHORT and TCGA-LAML), respectively. The datasets used in the manuscript are described in **Supplementary Table 3**.

### *In-vivo* AML murine models and bone marrow analysis

Animal studies were approved by Institutional Committee for Animal Welfare and by the Italian Ministry of Health and performed in accordance with national law D.lgs 26/2014 (authorization n. 781/2018-PR). For the experiments involving C1498 intra-bone (i.b.) injection, at day 0, 2×10^5^ Scr-C1498 or shZeb1-C1498 cells were injected into the tibia of immunocompetent mice. After 34 days, mice were sacrificed.

For experiments using the IL-17A-neutralizing antibody, animals were injected with 5×10^5^ Zeb1^+^ or silenced C1498. After 3 days, they were randomized. IL-17A–neutralizing or isotype control Ab (50μg/mouse) were injected i.p. twice a week. Mice were sacrificed after 30 days and BM and liver were explanted for further FACS analyses.

For the intracellular staining, the Foxp3/Transcription Factor Staining Buffer Kit (Tonbo Biosciences) was used. Antibodies used for FACS analysis are listed in **Supplementary Table 1**. Samples were analyzed with the FACSCelesta flow cytometer equipped with FACSDiva software (v 6.0) (Becton Dickinson). Flow cytometry data analyses were performed using FlowJo software (v10.2).

#### Statistical Analysis

Statistical analyses have been performed with GraphPad Software (Prism 8). The statistic applied to every single experiment is shown in the relative figure legend. Parametric and non-parametric analysis (Student t test, Mann-Whitney test) have been applied according to data distribution. A one-way ANOVA analysis with Tukey’s or Dunnett’s multiple comparison has been applied according to multiple comparison.

#### Data availability

All data needed to evaluate the conclusions in the paper are present in the paper and/or the Supplementary Materials. Microarray results are available in the National Center for Biotechnology Information Gene Expression Omnibus repository (accession number GSE192473).

## RESULTS

### The median of *ZEB1* expression dichotomizes AML patients and defines patients with worse OS and peculiar immune features

To define the relevance of ZEB1 in AML we performed *in silico* analysis on 7 independent cohorts from publicly available datasets (GSE15434, GSE16015, GSE12417, GSE37642, GSE6891, GSE161532 and TCGA, **Supplementary Figure 1A**) for a total of 1325 AML patients. AML patients were subdivided into two groups considering the median value of *ZEB1* expression among patients to bisect *ZEB1*^high^ and *ZEB1*^low^ AML (**Figure 1A**). To identify the main characteristics of *ZEB1*^high^ and *ZEB1*^low^ AML we performed a pathway analysis (**Figure 1B**) pointing out differences in the expression of immune response programs, including inflammatory response, interferon gamma response, allograft rejection that were down-regulated in *ZEB1*^high^ patients. On the contrary *ZEB1*^high^ patients showed an enrichment in MYC, HEME metabolism, IL17A and TGFβ pathways. Interestingly among genes differentially expressed between *ZEB1*^high^ and *ZEB1*^low^ AML we found *MYC, SOCS2*, *ARGINASE 2* (*ARG2*) that were up-modulated in *ZEB1*^high^ patients (**Figure 1C**). Noteworthy, SOCS2 is relevant gene involved in controlling proliferation and stemness of HSC also marking unfavorable leukemia (Vitali et al., 2015a) and ARG2 was shown to be involved in the polarization of monocytes into an immune suppressive M2-like phenotype in the leukemia microenvironment (Mussai et al., 2013).

**Figure 1.**
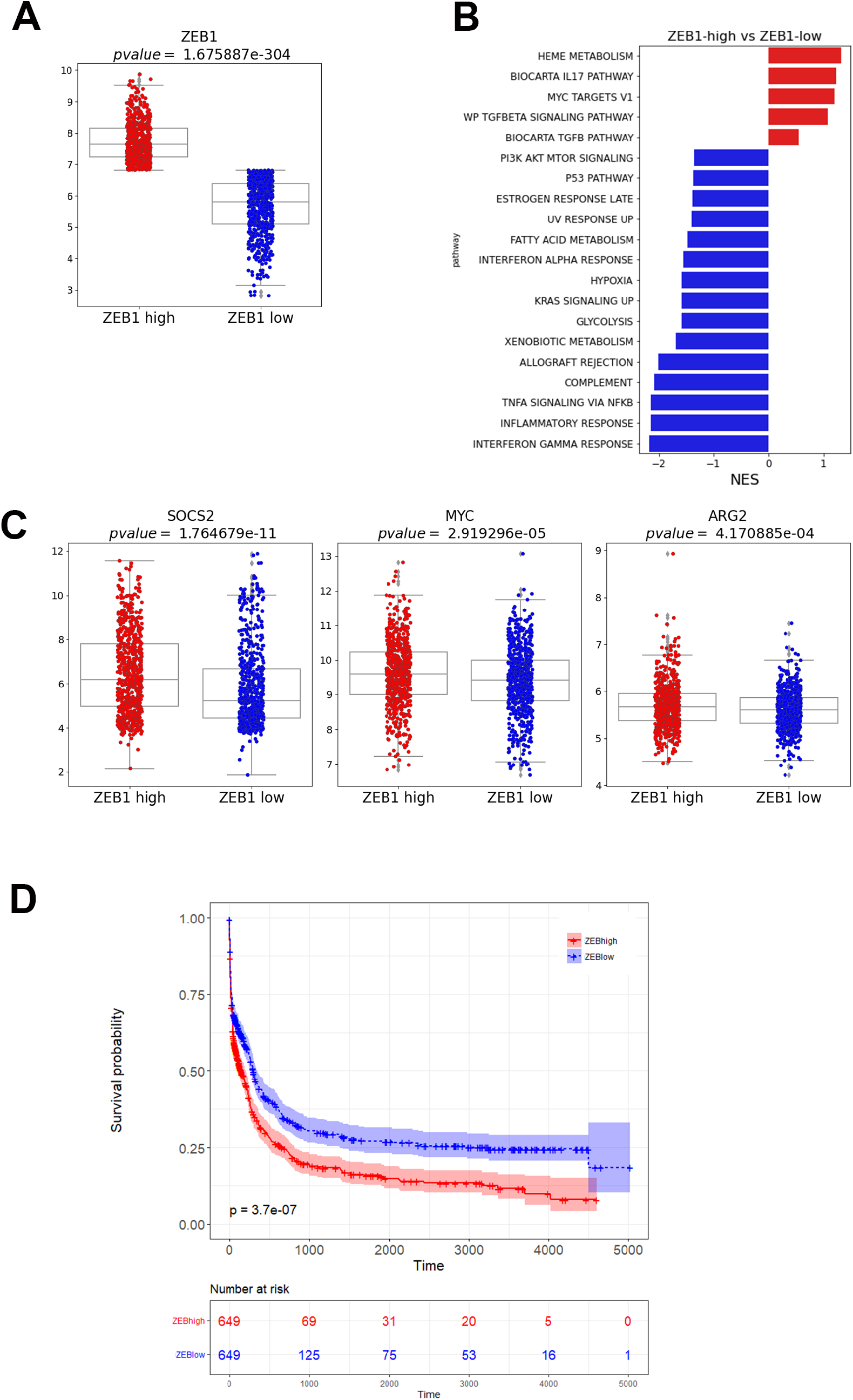
The median of *ZEB1* expression dichotomizes AML patients and defines patients with peculiar immune features and worse OS. **A.** *ZEB1* expression levels in the GEP analysis performed combining 7 independent AML cohorts (GSE15434, GSE16015, GSE12417, GSE37642, GSE6891, GSE161532 and TCGA) for a total of 1325 AML patients. **B.** Relevant pathways associated with *ZEB1*^high^ and *ZEB1*^low^ AML blasts in the 7 cohorts. **C.** Expression of selected differentially expressed genes (*MYC, SOCS2* and *ARG2*) in *ZEB1*^high^ and *ZEB1*^low^ AML patients. **D**. Kaplan-Mayer curves showing *ZEB1*^high^ and *ZEB1*^low^ patients overall survival (OS) combining the AML cohorts for which OS data were available (GSE12417, GSE37642, GSE6891, GSE161532 and TCGA for a total of 1298 patients.

Of note, combining datasets for which the overall survival was available (GSE12417, GSE37642, GSE6891, GSE161532 and TCGA) and for a total of 1298 patients, we found that ZEB1^high^ AML patients had a shorter OS than *ZEB1*^low^ patients (**Figure 1D**). This finding was also confirmed evaluating ZEB1 through IHC in a small separated cohort of AML patients (**Supplementary Figure 1B**).

Considering clinical features such as karyotype, French–American–British (FAB) classification and overall survival (OS) of AML patients, we observed a significant difference between *ZEB1*^high^ and *ZEB1*^low^ among cytogenetic subgroups (GSE6891, p=5.00E-07; TCGA-AML+ Beat AML, p=5.00E-07) (**Supplementary Figure 1C)**, with a higher and lower percentage of Normal Karyotype (NK)-and core-binding factor-mutated (CBF)-AML, respectively, in *ZEB1*^high^ versus *ZEB1*^low^ cases. The different distribution according to FAB classification was also confirmed (GSE6891, p=1.00E-07; GSE37642, p=4.60E-06; TCGA-AML+ Beat AML, p=3.00E-07) (**Supplementary Figure 1C)**, with *ZEB1*^high^ AML being enriched for more undifferentiated leukemia types. The large number of cases in the cohorts of public datasets allowed testing the association between *ZEB1* expression and the AML mutational profile. *ZEB1*^high^ and *ZEB1*^low^ cases showed a similar frequency of *ASXL1*, *DNMT3A*, *IDH1*, *IDH2*, *KRAS/NRAS*, *RUNX1*, *NPM1,* and *FLT3*-ITD mutations but differed for the concomitant presence of *FLT3*-ITD and *NPM1* mutations: 15.8% *vs.* 5.8% in *ZEB1*^high^ *vs. ZEB1*^low^, respectively (**Supplementary Figure 1D**). In line with cytogenetic data, *ZEB1*^high^ also included a higher percentage of *TP53*-mut patients (13.5% *vs.* 3.6% of ZEB1^low^, p=0.004), while *CEBPA* biallelic mutations were only present in the ZEB1^low^ cohort (6.6% vs. 0% of *ZEB1*^high^, p=0.003) (**Supplementary Figure 1D**).

The data as a whole suggests that ZEB1 might have an adverse impact on leukemia, potentially due to alterations in the immune microenvironment that can encourage immunosuppression.

### Zeb1 silencing impairs the invasive ability of leukemic C1498 cells without affecting their proliferation

To model the activity of ZEB1 in the context of leukemia, we started evaluating its expression in two widely used, and well-characterized, AML murine cell lines, C1498 and WEHI-3B cells (Mopin et al., 2016; Tomida, 1995). Western blot (WB) analysis (**Figure 2A**) and qPCR (**Figure 2B**) showed high levels of ZEB1, both at protein and mRNA levels, in C1498 cells and its paucity in WEHI-3B cells. Notably, the different expression of ZEB1 in C1498 *vs* WEHI-3B cells, correlated with the different expression of *Arg1* and *Tgfβ*, which were higher in C1498 AML cells and lower in WEHI-3B, respectively (**Figure 2C**).

**Figure 2.**
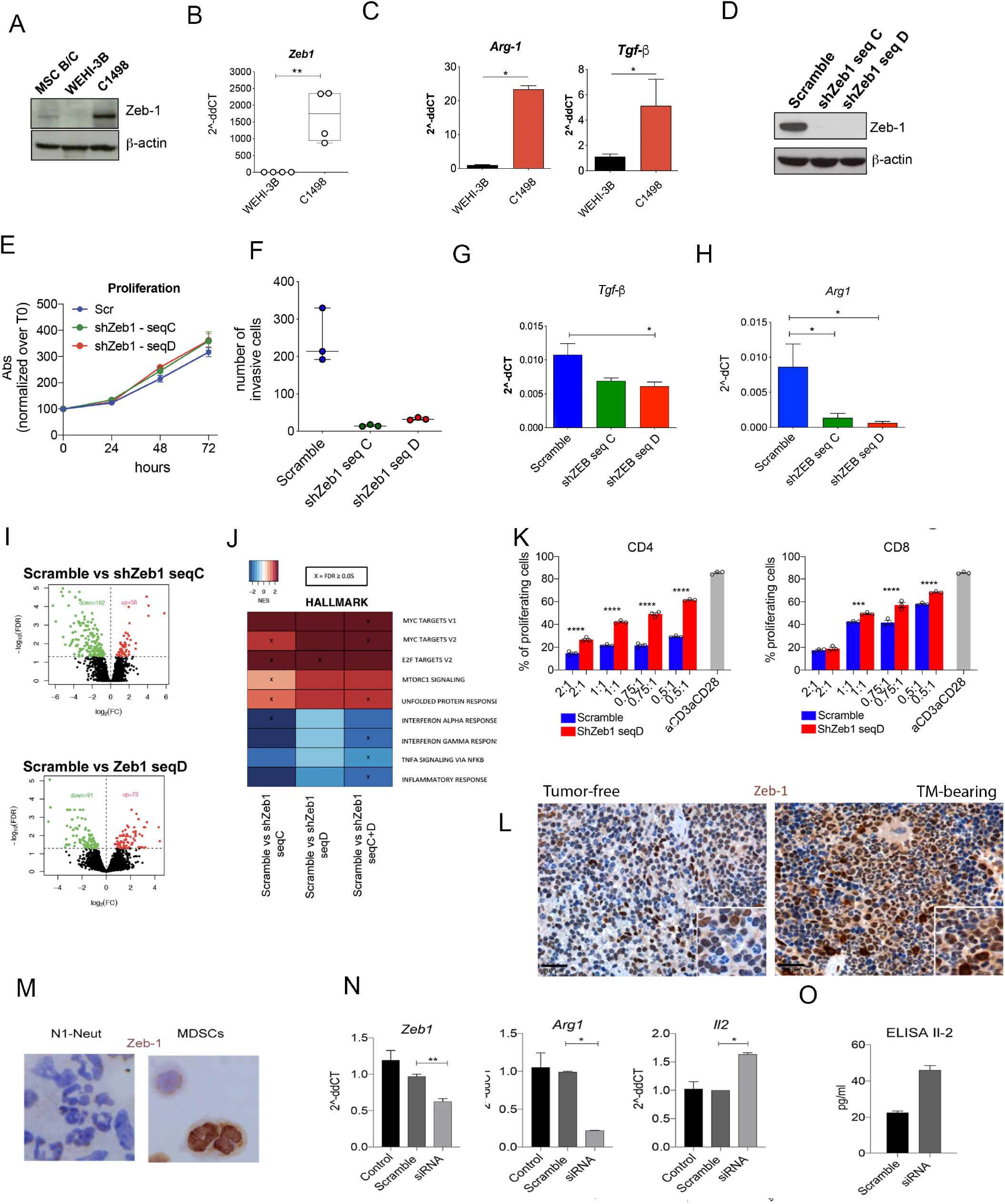
Zeb1 silencing impairs the invasive ability of leukemic C1498 cells without affecting their proliferation. **A.** Western blot analysis showing ZEB1 expression in WEHI-3B and C1498 murine AML cell lines. BM-derived mesenchymal stem cells (MSC) isolated from BALB/c (B/c) mice were used as ZEB1 positive control. β-actin was used as internal control. **B.** qPCR showing basal levels of *Zeb1* in WEHI-3B and C1498 murine AML cell lines. β-actin was used as internal control. (n=4, one experiment out of 2 performed, p=0.0050; Unpaired t test **p<0.01). **C.** qPCR showing the expression of *Arg1* and *Tgfβ* in C1498 and WEHI-3B cell lines. Results represent a pool of at least 2 independent experiments **D**. Stable knockdown of Zeb1 in C1498 using lentiviral vectors evaluated by Western Blot. β-actin was used as internal control **E.** XTT proliferation assay performed on Zeb1 expressing (Scr) and silenced cell lines. Cell proliferation for each time point was calculated as the (Absorbance (Abs) at 450nm – Abs at 670nm) t 24/48/72)/ (Abs at 450nm –Abs at 670nm) t 0 *100. Data represent a pool of 2 independent experiments; n=12/each experiment). **F.** *In-vitro* invasion assay using 24-well transwell plates (5 µm pore size) coated with 1 mg/ml Matrigel growth factor reduced basement membrane matrix. Five consecutive fields per transwell were counted. (n=3/each; one-way ANOVA; KW test p=0.0064). **G.** *Tgfβ* and **H.** *Arg1* expression levels in ZEB1 expressing (Scramble) and silenced cell lines. β-actin was used as internal control. **I.** Volcano plot showing the up-regulated and down-regulated genes in Scramble vs and silenced cells. **J.** Heatmap of canonical pathway enrichment analysis performed on *Zeb1* expressing vs silenced (sh*Zeb1*-seq C and sh*Zeb1*-seq D respectively) cells **K**. Suppression assay performed with Scramble and sh*Zeb1*-seq D silenced cells where irradiated C1498 cells were co-cultured with αCD3-αCD28 stimulated CFSE-labelled T-cells (total splenocytes) at different ratio. CD4^+^ and CD8^+^ proliferation was assessed 48h after. Data represent 1 experiment out of 3 performed (n=3/dilution; statistical analysis: unpaired t test ****p<0.0001). **L**. ZEB1 IHC staining performed on the spleens explanted from naïve or tumor bearing (breast cancer) mice. Original magnification, x200 and x630 (insets). Scale bar, 50 μm. **M.** ZEB1 IHC staining in MDSC isolated from mice bearing mammary tumors and N1 neutrophils isolated from agar plugs. **N.** *Zeb1* transient silencing (48h) in MDSCs sorted from tumor bearing mice. *Arg1* and *Il2* expression levels in Zeb1 silenced and Scramble control MDSCs. Untreated MDSCs were used as further negative control. Statistical analysis: One way ANOVA followed by Tukey’s multiple comparison. The p value relative to the comparison Scramble vs siRNA is shown in the figure (p=0.024 for *Arg1* and p=0.019 for *Il2*) **O.** Il2 levels detected by ELISA assay in Zeb1 silenced MDSC and Scramble control supernatants. (*p< 0.05; **p< 0.01; ***p<0.001; ****p<0.0001)

C1498 cells were stably silenced for *Zeb1* using a lentiviral vector also carrying a GFP tag to allow FACS sorting of infected C1498-GFP positive cells. All target-specific shRNA significantly decreased *Zeb1* expression compared to scramble transduced control (**Figure 2D**). In a reverse approach, we tried to overexpress *Zeb1* in WEHI-3B cells. Despite efficient transduction, according to GFP expression and *Zeb1* mRNA expression, (**Supplementary Figure 2A**) WEHI-3B cells failed to express ZEB1 protein (**Supplementary Figure 2B**). Similar result has been obtained transducing ZEB1 in human OCI-AML3 cells, in which mRNA but not ZEB1 protein was detectable upon gene transduction (not shown). Hence, we focused on murine and human cell lines endogenously producing ZEB1 and their silenced counterpart because it was unfeasible to force ZEB1 expression in low/negative cells.

Being ZEB1 associated with tumor aggressiveness, proliferation, and differentiation properties, we firstly investigated whether *Zeb1* silencing in C1498 cells could negatively impact cell growth or invasiveness. *In vitro* experiments showed that *Zeb1* down-regulation did not affect cell proliferation (**Figure 2E**) and only partially influenced C1498 differentiation state, inducing a slight increase in CD11b, Ly6C and Ly6G expression (**Supplementary Figure 2C**). Differently, *Zeb1* silencing significantly reduced the invasive properties of C1498 cells (**Figure 2F**), suggesting a possible role of ZEB1 in regulating the aggressiveness of AML cells. Interestingly, a relevant effect of *Zeb1* silencing in C1498 cells was the decreased expression of both *Tgfβ* and *Arg1* (**Figure 2 G-H**). Given the different behavior of *Zeb1*-expressing versus silenced C1498 cells, we compared the gene expression profiles of our cell lines. Results showed 56 up-regulated and 160 down-regulated genes in control versus sh*Zeb1*-seqC and 73 upregulated and 91 down-regulated genes in control versus sh*Zeb1*-seqD (**Figure 2I**). Notably, *Zeb1* expressing cells had higher expression of genes related to *Myc* and *E2f* pathways or involved in the mTOR signaling, compared with silenced counterpart (**Figure 2J**). Also, *Zeb1*-expressing cells had a decreased expression of inflammatory and immune-related pathways, among them *Tnf*, *Ifnα* and *Ifnγ* (**Figure 2J**). This data along with the down-modulation of *Arg1* and *Tgfβ* in *Zeb1* silenced cells, suggests a possible involvement of ZEB1 in immune regulation and particularly in immune suppression. To test the direct impact of ZEB1+ C1498 AML cells on T cell activity, we performed a suppression assay in which αCD3 αCD28 stimulated CFSE-labelled T cells were cultured with either *Zeb1*-silenced or *Zeb1*^+^ C1498 cells. Results show that *Zeb1*^+^cells more efficiently suppressed CD4^+^ and CD8^+^ T cell proliferation than sh*Zeb1*-seqD-silenced cells (**Figure 2K**). Notably, the link between ZEB1 and immunosuppression via Arginase I regulation was confirmed in myeloid derived suppressor cells (MDSCs), a subset of myeloid cells that expand in solid tumors to mediate immune suppression (van Vlerken-Ysla et al., 2023).Using a mammary tumor model previously characterized for the capacity to expand MDSCs (Sangaletti et al., 2016)we show that *Zeb1* is highly expressed by MDSCs (**Figure 2L**). ZEB1 protein level was higher in MDSCs isolated from the spleen of tumor-bearing mice than in inflammatory neutrophils (Sangaletti et al., 2012) (**Figure 1M**). Notably, transient knockdown of *Zeb1* in MDSCs (**Figure 2N**) was associated with decreased expression of *Arg1* paralleled by increased *Il2* (**Figure 2N**), a T cell activating cytokine, at both transcriptional and protein levels (ELISA, **Figure 2O**).

### Impaired BM engraftment of Zeb1-silenced AML cells is associated with cytotoxic CD8^+^ T cells expansion

To test the immune regulatory activity of ZEB1 *in vivo*, *Zeb1*-expressing (scramble transduced) or -silenced (sh*Zeb1*) C1498 cells were injected orthotopically into the tibia of immunocompetent, syngeneic C57BL/6 mice to better recapitulate the AML microenvironment within the BM. Lower frequency of GFP^+^ leukemic cells was found in the BM of mice injected with sh*Zeb* versus scramble transduced C1498 cells (**Figure 3A** for sh*Zeb1*-seqD **and Supplementary Figure 3A** sh*Zeb1*-seqC). This finding was associated with reduced extent of diffuse and nodular blast infiltration of liver parenchyma occurring in silenced versus control injected groups, (**Figure 3B**). Notably, the silenced group, exemplified by sh*Zeb1*-seqD cells, showed leukemic cells mostly confined around blood vessels, suggesting a limited invasiveness of silenced than control cells (**Figure 3C**). A reduction in C1498 take was also observed in the ovary of in mice injected with sh*Zeb1-cells* (**Supplemental Figure 3B**).

**Figure 3.**
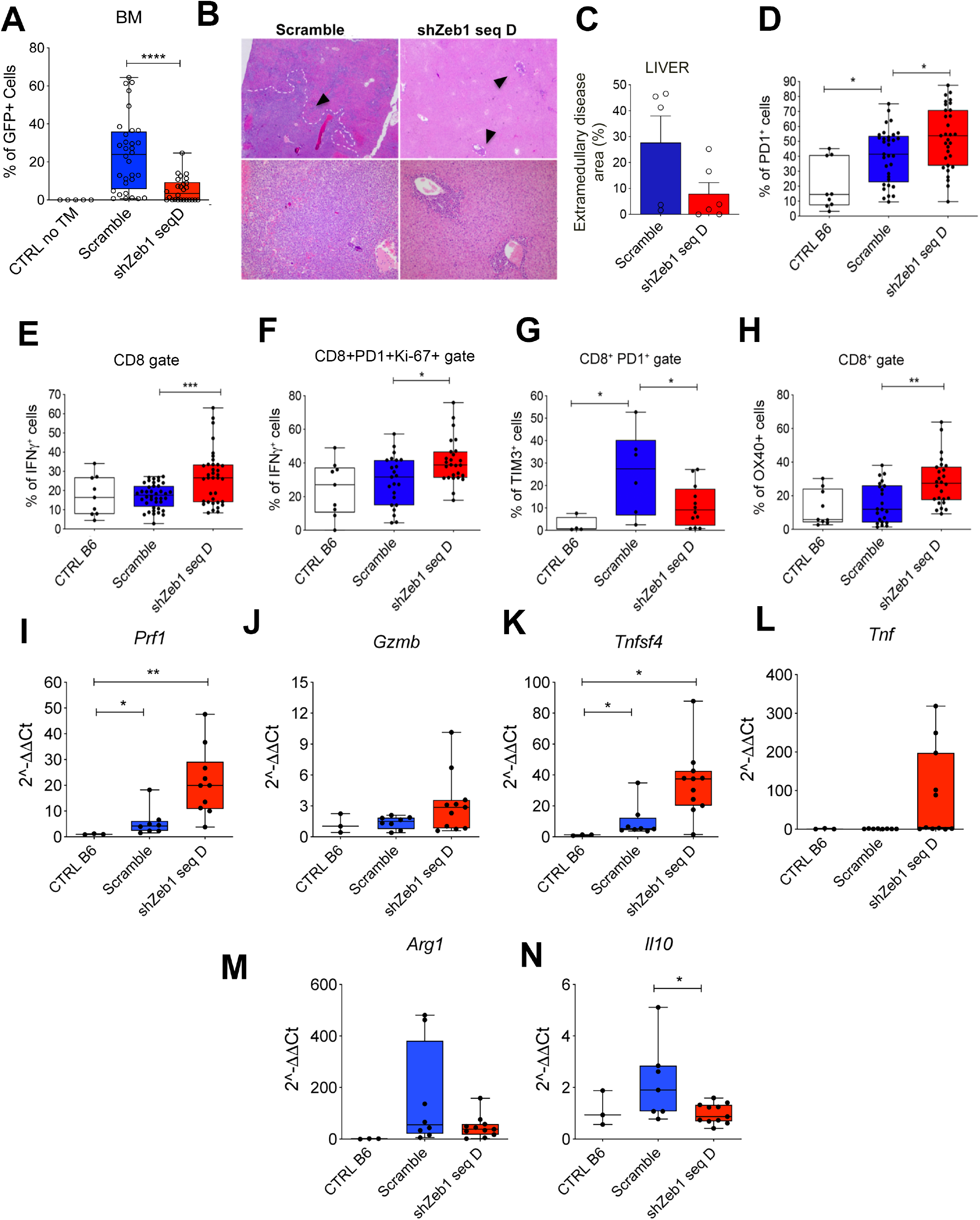
Zeb1 down-regulation was associated with the promotion of a cytotoxic microenvironment. **A.** Scramble (n=32) and sh*Zeb1*-seq D (n=26) were injected in the tibia of immunocompetent C57BL/6 mice and the percentage of GFP+ cells within the BM was evaluated with flow cytometry 34days post injection. Naïve mice (n=5) were used as controls. Data represent a pool of 3 independent experiments (Statistic: One-way ANOVA; for multiple comparison ****p<0.0001; KW test p<0.0001). **B.** Representative hematoxylin and eosin staining of liver explanted from Scramble-injected and sh*Zeb1*-seq D injected mice. 10X and 20X magnifications are shown. **C.** Extramedullary disease area quantification of liver explanted by Scramble-injected and sh*Zeb1*-seq D injected mice. Data represent the quantification of an individual experiment (n=5; unpaired t test; p=0.09). **D.** Frequencies of CD8^+^PD1^+^ within the BM of control mice (CTRLB6, n=9) or mice injected with *Zeb1*-expressing (Scramble, n=37) or silenced cells (sh*Zeb1-*seqD, n=31). Data represent a pool of 3 independent experiments (Statistical Analysis: One-way ANOVA followed by Tukey’s multiple comparison, p=0.0001; the p values relative to the comparisons Scramble vs CTRL or vs shZeb1 seq D are shown in the figure; *p<0.05). **E.** Frequencies of CD8^+^IFN-γ^+^ within the BM of control mice (CTRL B6, n=9) or mice injected with Zeb1-expressing (Scramble, n=39) or silenced cells (sh*Zeb1*-seqD, n=35). Data represent a pool of 3 independent experiments, (Statistical Analysis: one-way ANOVA followed by Tukey’s multiple comparison, p=0.0004; the p value relative to the comparisons Scramble vs shZeb1 seq D are shown in the figure; ***p<0.001). **F.** Frequencies of total CD8^+^PD1^+^Ki67^+^IFN-γ^+^ cells within the BM of control mice (CTRL B6, n=9) or mice injected with Zeb1-expressing (Scramble, n=39) or silenced cells (sh*Zeb1*-seqD, n=23). Data represent a pool of 2 independent experiments. (statistical Analysis: one-way ANOVA followed by Tukey’s multiple comparison, p=0.0038; the p value relative to the comparisons Scramble vs shZeb1 seq D is shown in the figure; *p<0.05). **G.** Frequencies of CD8 exhausted (PD1^+^TIM3^+^) cells within the BM of control mice (CTRLB6, n=5) or mice injected with Zeb1-expressing (Scramble, n=6) or silenced cells (sh*Zeb1*-seqD, n=12). statistical Analysis: one-way ANOVA followed by Tukey’s multiple comparison, p=0.0178; the p value relative to the comparisons Scramble vs CTRL or Zeb D is shown in the figure; *p<0.05 **H**. Percentage of CD8^+^OX40^+^lymphocytes within the BM of mice (CTRL B6, n=9, Scramble, n=22, sh*Zeb1*-seq D, n=26). Data represent a pool of 2 independent experiments (statistical Analysis: one-way ANOVA followed by Tukey’s multiple comparison, p=0.0009; the p value relative to the comparisons Scramble vs shZeb1 seq D is shown in the figure; **p<0.01). mRNA levels of **I.** *Perforin1* (*Prf1*) (KW test, p=0.001; *p<0.05); **J.** *Granzyme B* (*Gzmb*); **K.** *Ox40l* (*Tnfsf4*) (KW test, p=0.002; **p<0.001) and **L.** *Tnf*. **M.** *Arginase 1* (*Arg1*) (KW test, p=0.0133; *p<0.05) and **N.** *Il10* (KW test, p=0.0458; *p<0.05) within the BM of mice injected with C1498 expressing (n= 8) or silenced for Zeb1 (n=11). BM of naïve mice (n=3) was used as control.

The relevance of *Zeb*1 in shaping the AML immune microenvironment was investigated by multiparametric flow cytometry analysis performed on BM cells of mice injected *i.b.* with sh*Zeb1*-silenced or Zeb1+ C1498 AML cells.

We found increased frequency of CD3+ cells in the BM of mice injected with *Zeb1*-silenced than *Zeb1*+cells (**Supplementary Figure 3C**). This increase in T cells was not due to major CD4+ cell increment (**Supplementary Figure 3D)**, but mainly to a significant expansion of CD8+ T cells in the BM of mice bearing sh*Zeb1*-seq-D cells versus scramble treated C1498 cells (**Supplementary Figure 2E-F**).

Looking at the CD8 subpopulations, we observed a higher frequency of CD8^+^PD1^+^ (**Figure 3D**) and CD8^+^IFNγ^+^ (**Figure 3E**) T cells in the BM of mice receiving *Zeb1*-silenced than *Zeb1*+ control cells. Since PD1 is expressed on activated CD8 T-cells but it is also a marker of exhaustion, another marker of exhaustion, such as TIM3, was evaluated along with the ability to produce IFNγ. We found increased CD8^+^PD^−^1^+^Ki67^+^IFNγ^+^ cell fraction (**Figure 3F and Supplementary Figure 3G**). Concomitantly, the reduction of PD1^+^TIM3^+^ cell frequency (**Figure 3G and Supplementary Figure 3H**) in the BM of mice injected with *Zeb1* silenced cells confirms that *Zeb1* down-regulation in leukemic cells unleashes the expansion of activated T cells. Accordingly, the frequency of OX40+CD8+ lymphocytes was higher in BM injected with *Zeb1*-silenced cells than controls (**Figure 3H**).

The activation of CD8^+^ T cell in the BM was supported by qPCR analysis performed on total BM cells showing the up-regulation of *Perforin1 (Prf1)* and *Ox4Ol (tnfsf4)* and a trend in increase of *Granzyme B* and *Tnf* (**Figure 3I-L**) in mice injected with sh*Zeb1*-seq D compared with controls. This phenotype was paralleled by the decrease of genes encoding for immunosuppressive molecules, including *Arg1* and *Il10* (**Figure 3M-N**).

### IL17A is responsible for ZEB1-driven extramedullary infiltration of leukemia cell *in-vivo*

To further characterize ZEB1-dependent immunosuppressive effect on the BM microenvironment, we also analyzed the CD4^+^ population in the BM of mice injected with sh*Zeb1*-silenced or scramble transduced C1498 AML cells. Although the overall frequency of CD4 T cells was non different, the fraction of Treg cells was slightly reduced (**Figure 4A**) in favor of a statistically significant expansion of TNF-producing CD4^+^ cells (**Figure 4B**) and a trend toward increase of IFN-γ^+^ CD4^+^ (**Figure 4C**) cells in the *Zeb1*-silenced group. Furthermore, an expansion of CD3^+^IL-17^+^, Th17, and Treg IL-17^+^ T cells occurred in the BM of mice injected with ZEB1+ control cells in comparison to *Zeb1* silenced AML cells or normal mice (**Figure 4D-F and Supplementary Figure 3I-J**).

**Figure 4.-.**
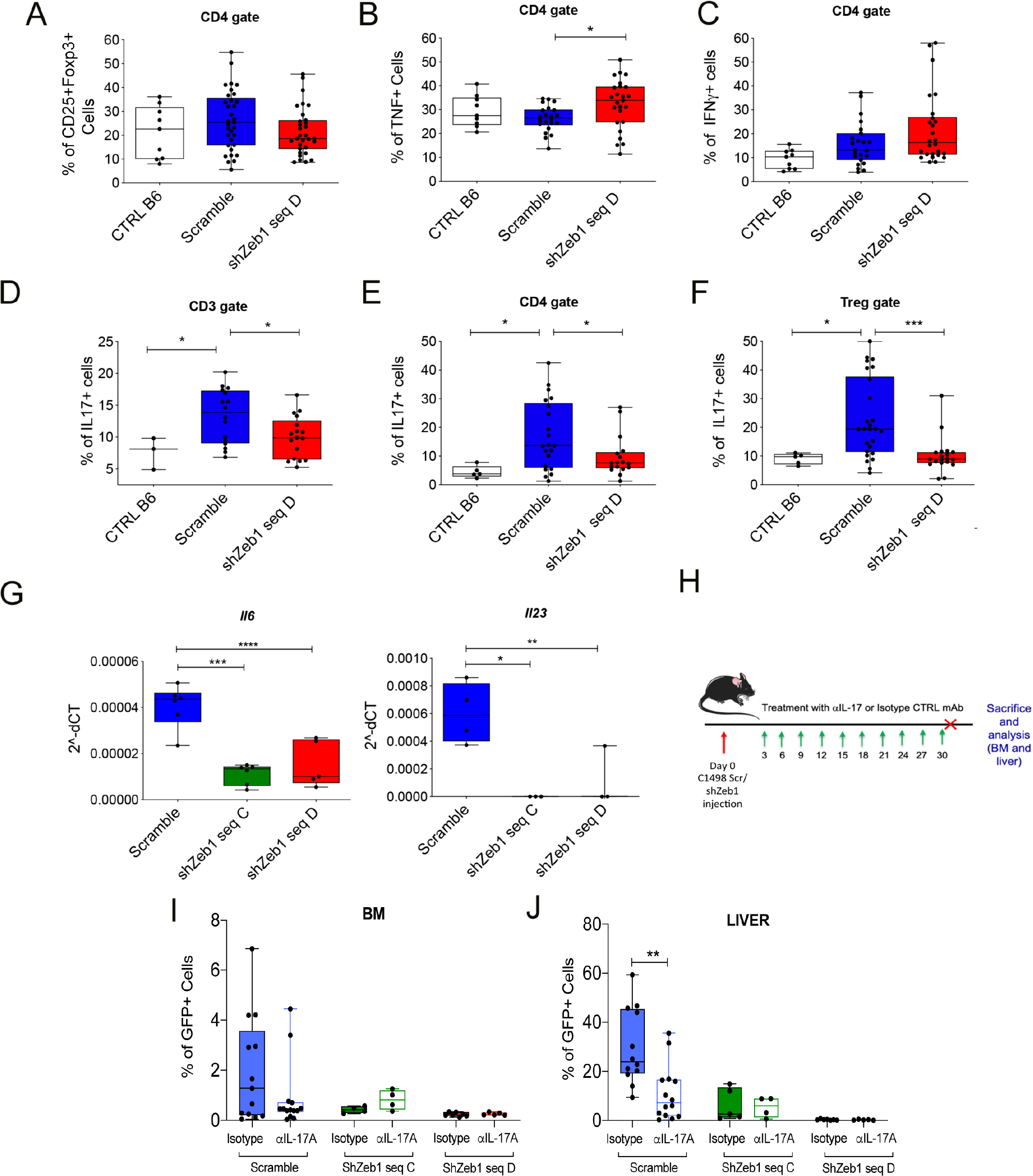
Zeb1 expression is associated with an expansion of lymphocytes producing IL-17A that in turn promotes AML aggressiveness. **A.** Frequency of Treg (CD4^+^CD25^+^Foxp3^+^) within the BM of control mice (CTRL B6, n=9) or mice injected with ZEB1-expressing (Scramble, n=31) or silenced cells (sh*Zeb1*-seqD, n=28). Data represent a pool of 2 independent experiments. Frequency of activated CD4 lymphocytes producing B. TNF and C. IFN-γ (CTRL B6, n=9, Scramble, n=21, sh*Zeb1*-seq D, n=25). Data represent a pool of 2 independent experiments Statistic: KW test, p=0.0388; For the multiple comparisons: *p<0.05; D. Frequency of IL-17^+^ CD3^+^ cells (CTRL B6, n=3, Scramble, n=16, sh*Zeb1*-seq D, n=17), Statistic: KW test, p=0.0289; For the multiple comparisons: *p<0.05. E. IL-17^+^ CD4^+^ cells (CTRL B6, n=5, Scramble, n=21, sh*Zeb1*-seq D, n=17; Statistic: KW test, p=0.0115; For the multiple comparisons: *p<0.05) and of F. Treg (CD4^+^CD25^+^Foxp3^+^) producing IL-17A (CTRL B6, n=5, Scramble, n=25, sh*Zeb1*-seq D, n=19). Statistic: KW test, p=0.0005; For the multiple comparisons: *p<0.05; ***p<0.001. G. *Il6* (statistic: ANOVA, Tukey’s multiple comparison test; ***p<0.001;****p<0.0001), and *Il23* (statistic: ANOVA, Tukey’s multiple comparison test; *p<0.05;**p<0.01) expression levels in silenced cells and Scramble. H. Immunocompetent mice were injected with either C1498 Scramble or Zeb1 silenced cells (5×10^5^ cells i.v.) at day 0 and then treated with anti-IL17 neutralizing antibody or its isotype control (50 μg. i.p.) every 3 days. Mice were sacrificed after 30 days and BM and livers were harvested for FACS analysis. Frequency of GFP+ cells within the I. BM and J. liver of mice injected with Zeb1 expressing or silenced cells and treated with Isotype control (Scramble n=12, shZeb1 seq C n=4-5, shZeb1 seq D n=7) or αIL-17A (Scramble n=12, shZeb1 seq C n=4-5, shZeb1 seq D n=7) Statistic: Mann-Whitney t test **p<0.01; p=0.0011.

This data might be explained by the significant down-modulation of *Il6* and *Il23A* in *Zeb1*-silenced cells (**Figure 4G**), which together with *Tgfβ* are the main regulator of Th17 differentiation (Geginat et al., 2016).To test the relevance of IL-17A production by CD3^+^, CD4^+^, and Tregs on AML growth and dissemination *in-vivo*, IL-17A neutralizing or isotype-matching mAbs were given every 3 days to mice implanted with *Zeb1* competent and silenced cells (**Figure 4H**).

In mice injected iv, IL-17A blockade slightly reduced leukemic cell take in the BM (**Figure 4I**) but significantly impaired the infiltration of C1498 cells to the liver compared to mice treated with the isotype control only in Scramble injected mice (**Figure 4J**). The almost unchanged engraftment of C1498-GFP cells in presence of a strong reduction in liver spreading under treatment with anti-IL-17A-neutralizing Abs, suggests a major role of IL-17A in AML dissemination.

To better understand how IL-17A promotes AML dissemination in relation to ZEB1 expression, *Zeb1*-silenced and *Zeb1*+ C1498 cells were stimulated with mrIL-17, and tested for cell proliferation. Interestingly, rIL17 significantly increased the proliferation of *Zeb1*^+^ cells (**Figure 5A**) and the expression of *Socs2,* which of note was one of the relevant gene differentially expressed among *ZEB1*^high^ and *ZEB1*^low^ AML patients, and *mmp9* the latter is relevant marker of AML (Pirillo et al., 2022) also directly associated to blast invasion (Feng et al., 2011) (**Figure 5B**). In addition, rIL-17 exposure also induced *Il6* and *Tgfβ* expression (**Figure 5C**), thus supporting the idea that AML cells can self-activate a Th17-maintaining loop. Notably, the expression of IL17A receptor remained unchanged in *Zeb1*-silenced and Zeb+ cells (**Supplementary Figure 4A**).

**Figure 5.-.**
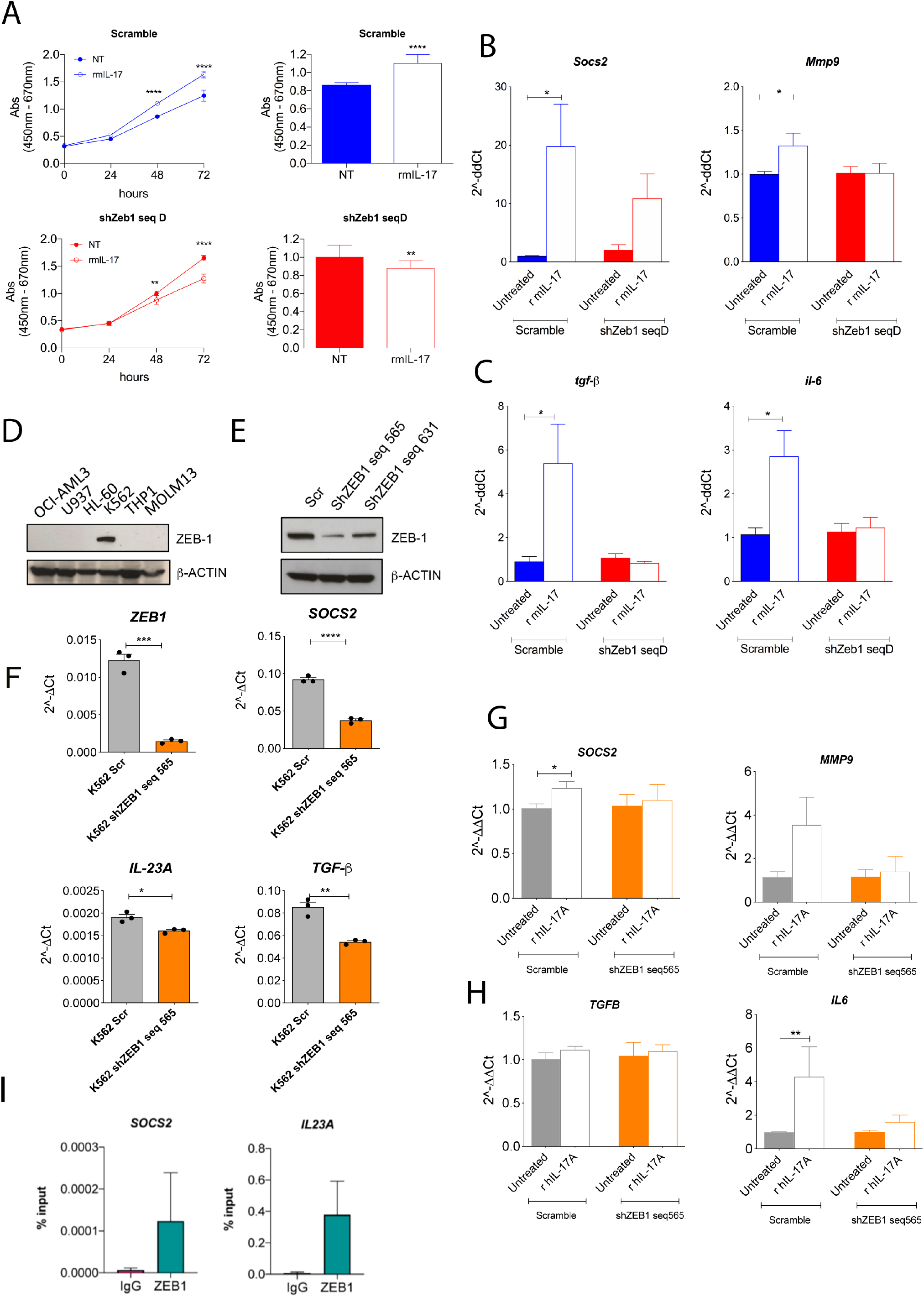
IL-17A stimulation promotes the expression of genes associated to leukemic cell aggressiveness. **A.** Cell proliferation of C1498 expressing (scramble) or silenced (sh*Zeb1*-seq D) upon IL-17A stimulation (50 ng/mL) assessed by Xtt assay after 24h, 48h and 72h. Cell proliferation for each time point was calculated as the (Absorbance (Abs) at 450 nm – Abs at 670 nm). Data represent a pool of 2 experiments (Two-way ANOVA, Multiple comparison test: ** p< 0.01;****p<0.0001). **B.** qPCR showing the induction of *Socs2*, *Mmp9* and **C.** *Tgfβ and Il6* upon stimulation (7 days) with IL-17A 50 ng/mL in Zeb1 expressing cells. Statistic: Paired *t* test; p=0.0255) **D.** ZEB1 expression in a panel of 6 different leukemic cell lines evaluated by Western Blot. **E.** Stable knockdown of ZEB1 in K562 using lentiviral vectors evaluated by Western Blot. β-actin was used as internal control. **F.** *ZEB1* (p=0.0002), *SOCS2* (p<0.0001)*, IL23A* (p=0.0125), and *TGFβ* (p=0.0025) expression levels in silenced cells and Scramble. Data represent a pool of 3 experiments (Statistics: Unpaired t test) **G.** qPCR showing the induction of *SOCS2* (*p< 0.05), *MMP9* and **H.** *TGF-β, IL6* (**p< 0.01) upon stimulation (7 days) with IL-17A 50 ng/mL in ZEB1 expressing cells.; **I.** Chromatin immunoprecipitation assay of ZEB1 showing the direct binding to SOCS2 and IL23A in K562 cells. The fraction of chromatin bound to the promoter, with IgG negative control or anti-ZEB1 antibody is represented as a percentage of input. Data represent a pool of 2 independent experiments.

To corroborate our mouse data in the human setting, we initially investigated ZEB1 expression in a panel of 6 different leukemic cell lines. Among them, only the K562 was expressing ZEB1 (**Figure 5D**) allowing the generation of its *ZEB1*-silenced counterpart (**Figure 5E**). Although without differences in the surface markers (**Supplementary Figure 4B**), the *ZEB1*-silenced variant showed reduced expression of *SOCS2, TGFβ* and of IL23, a cytokine involved in Th17 maintenance (**Figure 5F**) (Stritesky et al., 2008). As for murine AML cells, rhIL17 stimulates K562 cell proliferation (**Supplementary Figure 4C**) in association with up-regulation of *SOCS2*, *MMP9*, and *IL6,* but not *TGFβ* (**Figure 5G, H**).

To identify the direct targets of ZEB1 among molecules affected during its silencing, we performed Chromatin immunoprecipitation (ChIP) experiment followed by a qPCR on the obtained DNA. With this experiment, in human K562 cells, we were able to validate the binding of ZEB1 to the promoter of *SOCS2* and *IL23A* (**Figure 5I**). Notably, we further confirmed these data also analyzing publicly available Chip-seq data, which although performed on lymphoma cells (GM12878), confirmed the direct binding of ZEB1 to the promoter of *SOCS2* and *IL23A* (**Supplementary Figure 4D**).

To further challenge the association between ZEB1 expression in leukemic blasts and Th17, we performed a double IF staining on archival BM biopsies from 26 AML patients divided into ZEB1^high^ and ZEB1^low^ according to the median number of ZEB1+ nuclei, which ranged from 0.5 up to 60% (**Figure 6A**). Notably, we found a higher number of CD3^+^IL-17^+^ cells in ZEB1^high^ than ZEB1^low^ AML cases (**Figure 6B, C**) with a positive correlation between ZEB1 levels and the number of CD3^+^IL-17A^+^ cells (**Figure 6D**). Accordingly, a gene signature able to identify Th17cells in TSGA AML cohorts using the GEPIA2 tool revealed an association with poor outcome in terms of overall survival (**Figure 6E**) in patients enriched for this signature.

**Figure 6.-.**
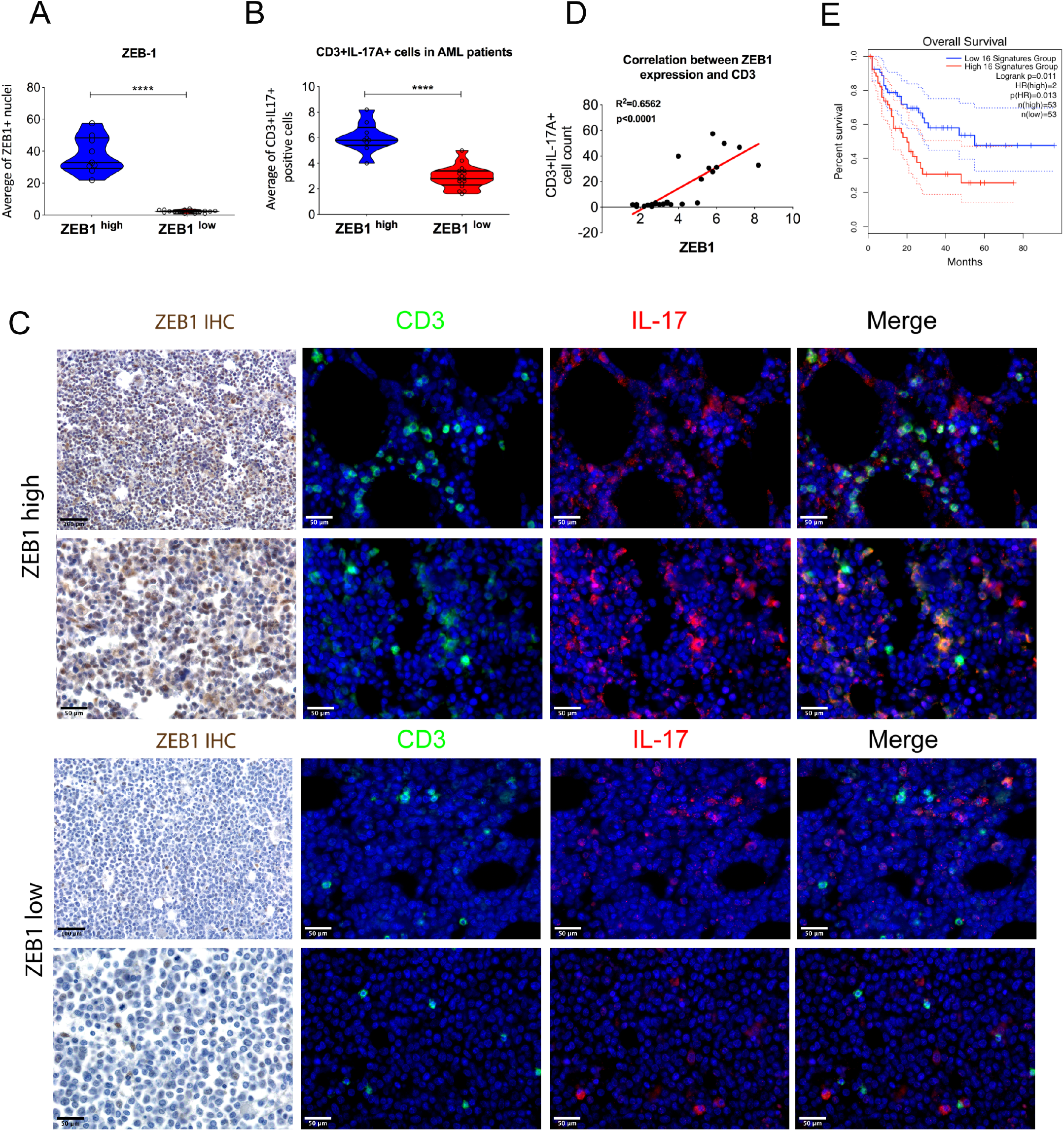
ZEB1^high^ levels in AML patients positively correlates with the expansion of IL17+ CD3 cells. **A**. ZEB1 quantitative analysis of immunohistochemical staining performed using the Image Analysis software provided by Leica (“nuclear hub” tool). (Statistic: Mann-Whitney *t* test****p< 0.0001) **B.** Quantification of CD3+IL-17+ cells within the BM of ZEB1^high^ (n=9) and ZEB1^low^ (n=16) AML patients. (Statistic: Mann-Whitney *t* test****p< 0.0001) **C.** Representative immunohistochemical staining for ZEB1 and immunofluorescence for CD3+IL-17A+ evaluation (CD3 in green and IL-17A in red) performed on 26 archival BM biopsy of AML patients (University of Palermo cohort). Original magnification, x200 and x400. Scale bars, 50 and 100 μm. **D.** Positive correlation between ZEB1 positive cells and CD3+IL-17A+ infiltrate. Statistics: Pearson’ correlation; R= 0.6265; p< 0.0001. **E.** Overall survival of AML patients (TCGA) with high and low Th17 infiltration. 16 genes (*CXCL3, IL22, IL3, CCL4, GZMB, LRMP, CCL5, CASP1, CSF2, CCL3, TBX21, ICOS, IL7R, STAT4, LGALS3* and *LAG3*) were used to define specific cell populations.

### Relevance of Th17-ZEB1 axis in AML relapse

Given the known activities of ZEB1 in drug resistance (Meidhof et al., 2015) we investigated its possible role in AML relapse. We performed in silico studies using gene expression profiling (GEP) analysis on paired AML blasts obtained at diagnosis and relapse (GSE66525) (Hackl *et al*., 2015). We found a further increase in ZEB1 expression in relapsed samples (**Figure 7A**), along with a trend of increased MMP2 and SOCS2 expression (**Figure 7B-C**). Interestingly, in the same patients, the higher ZEB1 levels in relapsed patients were associated with the enrichment in the Th17 pathway (**Figure 7D**). Moreover, investigating the differentially expressed hallmarks at relapse compared with diagnosis, we found an up-regulation of MYC targets and UV response, and a downregulation of inflammatory response, allograft rejection, and interferon gamma response (**Figure 7E**), all pathways related to ZEB1-positive blasts. This finding supports the hypothesis that relapsed AML is enriched in ZEB1-positive blasts and maintains the ZEB1-driven Th17 skewing.

**Figure 7.**
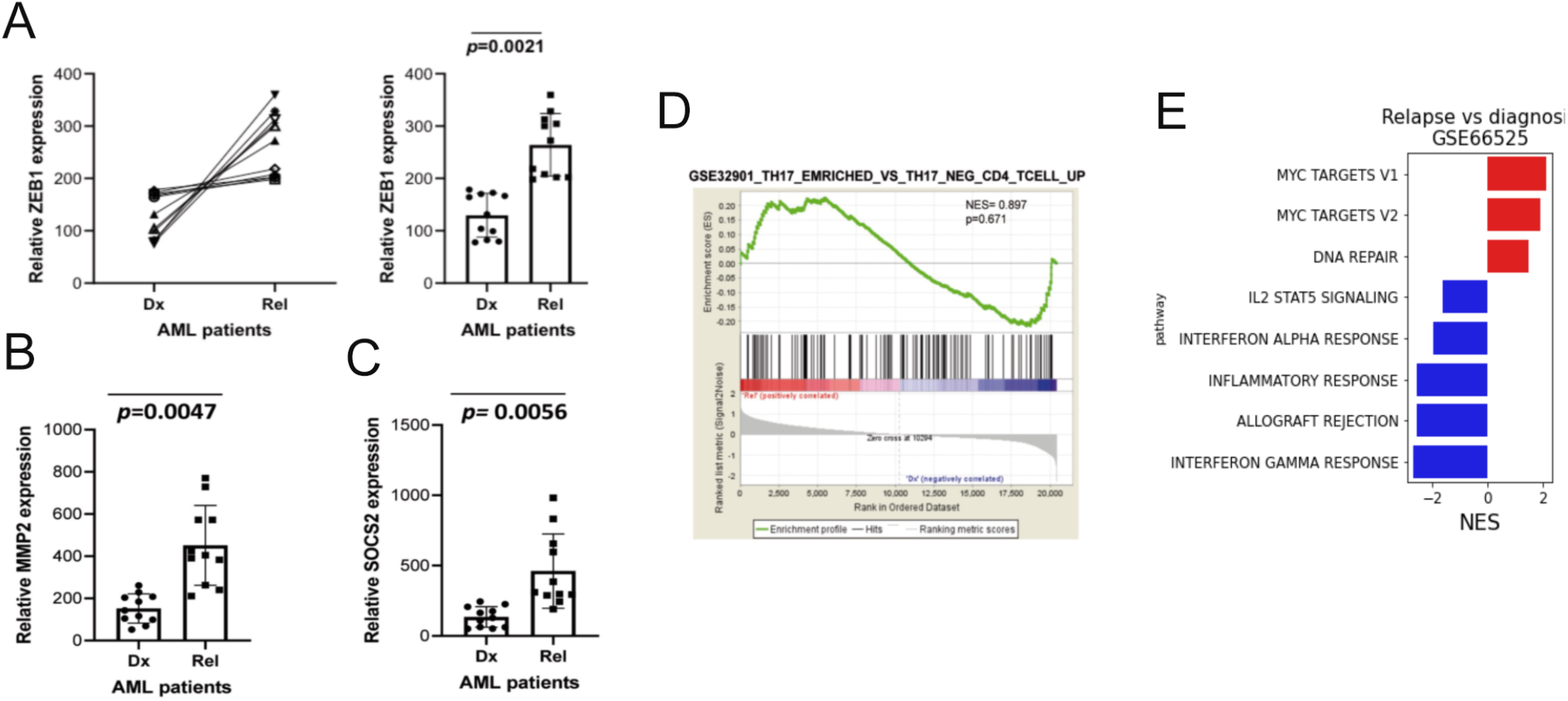
Relevance of Th17-ZEB1 axis in AML relapse. **A.** *ZEB1* expression levels at diagnosis and relapse in 11 AML patients (GSE66525). Expression levels of **B.** *MMP2* and ***C.*** *SOCS2* at diagnosis and relapse in GSE66525 **D.** IL-17 enrichment pathway at relapse vs at diagnosis. **E.** hallmark pathways enriched or downregulated in relapsed patients compared with patients at diagnosis.

## Discussion

Nowadays it is widely accepted that AML cells can influence the BM microenvironment to their own advantage such to create a peculiar niche that supports their survival, resistance to therapy and immune evasion. Nevertheless, beside IFNγ, no other molecular drivers active in molding the BM immune microenvironment under AML influence have been characterized in depth (Corradi et al., 2022; Vadakekolathu *et al*., 2020). Here, we demonstrate a formerly undiscovered ability of the EMT-regulator ZEB1 to shape the BM immune microenvironment, when expressed by leukemic blasts, sustaining AML progression. Our data indicate that ZEB1 directly orchestrates T–cell suppression via Arginase, and promotes Th17 expansion; the combination of these features negatively affects patients’ OS.

Previous studies performed on AML characterized by the *MLL-AF9* fusion gene, showed ZEB1 expression associated with aggressive LT-HSC-derived AML, and with reduced OS. Regarding the potential oncogenic activity of ZEB1 in AML (Li *et al*., 2021) (Shousha *et al*., 2019) (Stavropoulou *et al*., 2016) discordant evidences have been published. Almotiri and coll. postulated that ZEB1 acts as a transcriptional regulator of haematopoiesis and that its expression is required to suppress leukemic potential in AML models (Almotiri et al., 2021). This data might fit with our inability of over-expressing ZEB1 protein despite efficient gene transduction and mRNA expression in both moue and human cell lines that endogenously are low/negative for Zeb1. Differently, the possibility of silencing ZEB1 where it is spontaneously expressed suggest a cell-dependent protein dosage limitation. Our analysis of *ZEB1* expression and distribution in larger datasets, including those evaluated by Almotiri and coll., showed that not all AML patients express *ZEB1* at level lower than healthy controls. Rather, a fraction of them expresses significantly higher levels of *ZEB1* than controls, particularly in the CN-AML cohorts. Also, our IHC/IF analyses confirmed higher inter-patient variability with some patients showing positivity for ZEB1 protein in almost all blasts and other patients showing ZEB1 expression confined to few blasts. To better dissect the biological and clinical features associated to *ZEB1*^high^ AML, we analyzed *in silico* data and dichotomized patients in two groups according to the median level of *ZEB1* expression that also bisects OS, FAB and karyotypes. At molecular level, *ZEB1*^high^ AML patients showed increased expression of pathways related to the Myc, SOCS2, IL-17, TGFβ and HEME metabolism and down-modulation of inflammatory pathways. The last combined to the up-regulation of ARG2 gene in *ZEB1*^high^ AML have suggested the ZEB1 could entwines both pro-tumoral and immune suppressive features in AML that were accordingly demonstrated in immune competent mice.

In the C1498 AML mouse model, knock-down of *Zeb1* revealed that ZEB1+ leukemia cells may hinder the activity of CD4 and CD8 T-cells by directly modulating *Arg1* expression. Although the direct binding of ZEB1 to the *Arg1* promoter was not demonstrated, the observed decrease in *Arg1* expression resulting from transient silencing of MDSCs strongly supports this hypothesis. Additionally, in humans, the co-regulation of ZEB1 with ARG2 and the ENCODE project’s identification of ARG2 as one of ZEB1’s target genes further supports the notion that ZEB1 functions as a regulator of Arginase. Arginase 2 is a protein involved in several cellular functions, including polyamine synthesis and cellular energy metabolism. Although it is not completely elucidated whether it directly suppress immune function, its interaction with other molecules or cellular pathways can have indirect effects on immune function (Dowling et al., 2021; Grzywa et al., 2020). In acute myeloid leukemia (AML), an increased expression of Arginase 2 in blasts contributes to chemotherapy resistance suppressing immune responses by polarizing surrounding monocytes into a suppressive M2-like phenotype (Mussai *et al*., 2013). Therefore, whether Arginase 1 and 2 might play similar or divergent functions also according to their cellular source and species remains unclear. Interestingly, arginase 1 has been involved in the differentiation ofTh17 (Wu et al., 2016), which along with immune suppression represented the main features associated to ZEB1^high^ AML. It is noteworthy that, in both human and murine models, Th17 cells promoted the expression of SOCS2, a gene that is involved in the aggressiveness of leukemia (Vitali *et al*., 2015a; Vitali et al., 2015b). In this case, our Chip-qPCR analysis, along with data from the ENCODE project, demonstrated the direct binding of ZEB1 to the promoter of SOCS2 and IL23.

In humans, Th17 are prognostically relevant in AML and other cancers (Civini et al., 2013) (Han *et al*., 2014). Musuraca *and coll.* described a population of IL-17/IL-10-secreting immune suppressive Th17-cells that could identify AML patients with a higher risk of severe infections and relapse (Musuraca et al., 2015). The increase in ZEB1, along with Th17, SOCS2, and MMP2, in blasts at relapse compared to diagnosis, leads to the conclusion that ZEB1 expression in AML may be particularly relevant in identifying patients who are at a higher risk of relapse after chemotherapy and, more importantly, after allogeneic hematopoietic stem cell transplantation (HSCT). Even the improvement in the treatment, relapse still represents a common scenario in AML, occurring in 40–50% of younger and the great majority of elderly patients (Thol and Ganser, 2020).This relapse usually arises within the bone marrow, even if increasing reports highlighted the existence of extramedullary relapses, which involve skin and soft tissues (Harris et al., 2013).

Finally, our findings, linking ZEB1 to AML immune suppression, are mirrored in solid tumors where EMT factors contribute to immune evasion (Terry et al., 2017) (Dongre et al., 2017) (Plaschka et al., 2022), and highlights the need to further investigate the molecular mechanisms by which tumor intrinsic EMT-related pathways affect the microenvironment. The study suggests that EMT/ZEB1 could be a candidate predictive marker to be targeted using specific approaches.

## Acknowledgements

We thank Dr. Maria Teresa Majorini and Dr. Matteo Dugo for bioinformatic assistance and Matteo Milani for preliminary analysis. We thank Dr. Pagani M., Dr. Della Chiara G. and Dr. Bason R. for technical assistance with Chip-qPCR experiments. The authors also thank Ester Grande for providing administrative support. We thank HOVON for sharing data relative to the AML cohort included in the GSE6891.

## Author contributions

**B. Bassani**: performed cell culture, molecular experiments, flow cytometry experiments, statistical analysis and contributed to draft the manuscript. **V. Cancila** and **A. Gulino:** performed IHC experiments. **M. Ciciarello:** revised the manuscript. **C. Chiodoni** and **D. Lecis:** performed stable Zeb1 silencing and revised the manuscript. **G. Simonetti, A. Piva** and **E. Fonzi:** performed bioinformatics analysis. **L. Botti:** performed *in-vivo* experiments. **P. Portararo:** performed experiments on MDSCs and suppression assay. **M. Costanza:** performed ELISA assay for Il-2. **J. Schwaller**, **A. Tzankov**, **M. Ponzoni**, **F. Ciceri, N. Bolli** and **A. Curti** revised the manuscript. **A. Tzankov** also provided the independent cohort of 24 patients, performed IHC experiments and analyzed the results. **C. Tripodo**: supervised IHC experiments and revised the manuscript. **M.P. Colombo:** critically discussed data and revised the manuscript. **S. Sangaletti:** conceived the study, analyzed the data, performed statistical analysis and wrote the manuscript.

All authors read and approved the final manuscript.

