## Supplementary material for "ZEB1 shapes AML immunological niches suppressing CD8 T-cell activity while fostering Th17 cell expansion": Supplementary Methods Figures and Tables.pdf

#### Supplementary Methods

##### Cell cultures

The C1498 cell line, a murine AML cell line isolated from a leukemic 10-month-old C57BL/6 (H-2b) female mouse in 1941 205 was purchased from ATCC, while the WEHI-3B murine myelomonocyte cell line syngeneic to BALB/c mice was purchased from Sigma Aldrich (Merck, 86013003). K562 are a human erythroleukemia cell line isolated from the bone marrow of a 53-year-old patient. Cells were cultured in DMEM (Dulbecco's modified Eagle's medium) or RPMI-1640 (Thermo Fisher Scientific) supplemented with 10% fetal bovine serum (FBS; Thermo Fisher Scientific), 1% antibiotics (Thermo Fisher Scientific), 2mM glutamine, 1mM sodium pyruvate, 1mM HEPES and 1X Minimum Essential Medium (MEM) Non-Essential Amino Acids Solution, in a humidified atmosphere containing 5% $CO_2$ at 37°C. The neutralizing mouse IL-17A mAb (clone 17F3) was purchased from BioXCell.

#### *Quantitative immunolocalization analyses*

IHC analysis was performed on BM biopsies from 26 AML patients from the University of Palermo (Protocol 443/1/10/18, authorization number 09/2018). No clinical information concerning disease features and patients outcome are currently available.

For immunohistochemistry (IHC), human and murine bone marrow samples were fixed in 10% buffered formalin, decalcified using an EDTA-based buffer, and paraffin-embedded. Four micrometers tissue sections were deparaffinized and rehydrated. Novocastra Epitope Retrieval Solution (pH9) was used to unmask antigens in a thermostatic bath at 98°C for 30 min. Subsequently, the sections were brought to room temperature and washed in PBS. After neutralization of the endogenous peroxidases with 3% $H_2O_2$ and Fc-blocking by 0,4% casein in

PBS (Novocastra), the sections were incubated with primary antibodies listed in Table 3. IHC staining was developed using the Novolink Polymer Detection Systems (Novocastra) or IgG-Peroxidase specific secondary antibody (Sigma Aldrich) and DAB (3,3'-diaminobenzidine) as substrate chromogen. Anti-mouse and anti-goat (Alexa Fluor 488 and 568 conjugate) secondary antibodies were used for immunofluorescence (IF) and DAPI (4',6-diamidin-2-fenilindolo) for nuclei visualization. Slides were analyzed under a Zeiss Axioscope A1 microscope equipped with four fluorescence channels widefield IF. Microphotographs were collected using a Zeiss Axiocam 503 Color digital camera with the Zen 2.0 Software (Zeiss). Slide digitalization was performed using an Aperio CS2 digital scanner (Leica Biosystems) with the ImageScope software (Aperio ImageScope version 12.3.2.8013, Leica Biosystems). Quantitative analyses of IHC stainings were performed by calculating the average percentage of positive signals in five separate fields at medium-power magnification (X200) using the Nuclear Hub Image Analysis package and the result was expressed as a percentage.

##### *Extramedullary disease quantification*

Livers from i.b. injected mice either with Zeb1-expressing or -silenced C1498 cells were explanted after 34 days, washed in PBS and fixed in 10% neutral buffered formalin overnight before embedding in paraffin. Four-micrometers-thick tissue sections were deparaffinized using xylol and firstly rehydrated in 100% ethanol for 5 min. Then, sections were incubated in 95%, 80%, 50% ethanol for 5 min and finally washed in distillated water. Sections were incubated with hematoxylin for 8 min and then washed. Eosin was added to the tissue sections for 3 min and then washed. The stained sections were dehydrated in 70% ethanol for 2 min, 100% ethanol for 2 min and finally in xylol twice for 5 min. Sections were mounted using Eukitt (Biosigma). Infiltrated areas were then quantified using Leica software. For flow cytometry analysis, livers were mechanically smashed in DMEM with 10% FBS and then filtered through 70 μm cell strainer. Red blood cells were lysed using a solution of Ammonium-Chloride-Potassium lysing Buffer (ACK). If necessary a second step of filtration was made prior analysis by flow cytometry to avoid clogging issues.

##### *Immunoblotting*

40µg of the total protein lysate was separated on 8 or 12% SDS–polyacrylamide gel electrophoresis under reducing conditions and transferred onto nitrocellulose membranes (Amersham, Biosciences). Following blocking with 5% bovine serum albumin (BSA) and 0.1% Tween-20, the membranes were incubated with the antibodies listed in Supplementary Table 4 (1:1000 dilution) overnight at 4°C. After rinsing in tris-buffered saline (TBS) 0.1% Tween-20, membranes were incubated with horseradish peroxidase conjugated goat anti-rabbit secondary antibodies (Thermo Scientific; 1:2000) and reactions were visualized with the Western BLoT Quant HRP Substrate (TakaraBio).

##### *Invasion assay*

$2.5x10^5$ C1498 cells either with and without silenced *Zeb1* expression were resuspended in 200µL of serum-free high glucose DMEM and placed onto the upper chamber of a 24-well Transwell plate (5-µm pore size) coated with growth factor reduced matrigel (1mg/mL). 750µL of high glucose DMEM containing 10% FBS was added into the lower chamber. After 24h at 37°C and 5% $CO_2$, top chambers, containing non-migrated cells were removed, while cells that migrated into the lower chamber were counted using the all-in-one digital inverted fluorescence microscope (EVOS fl – advance microscopy group). 5 randomly selected fields per well were counted.

##### *Proliferation assay*

To assess the proliferation of Zeb1-expressing or -silenced cells, we used the colorimetric Xtt assay. This test is based on the cleavage of tetrazolium salts added to the culture medium and allow the evaluation of cell viability and proliferation. Briefly, $10^4$ cells were seeded in a 96-well plate in 100 µl of DMEM 10% FBS for four different time point, termed $t_0$, $t_{24}$, $t_{48}$ and $t_{72}$. For each time point, plated cells were incubated with 50µl XTT labelling mixture per well and incubate for 4h at 37°C and 5% $CO_2$. Absorbance of the formazan products was measured at 450nm Tecan's Spark Mircoplate reader, while the reference wavelength was read at 670nm.

#### *In vitro suppression assay*

4 $\times 10^5$ naïve C57Bl/6 splenocytes were labeled with CFSE (Carboxyfluorescein Succinimidyl ester; 10mM, SIGMA Aldrich) and co-cultured with irradiated (3Gy) C1498 Zeb1-expressing or -silenced cells at different ratio in presence of 2µg/ml of soluble anti-CD3 and 1µg/ml of anti-CD28 to activate lymphocytes. Each sample was seeded in triplicate. Proliferation of CD4 and CD8 T cells has been assessed after 48h by flow cytometry evaluating CFSE dilution in the $CD4^+$ and $CD8^+$ gated populations.

#### *MDSCs and neutrophil isolation from human and mice*

MDSCs ($CD11b^+Ly6G^+Ly6C^+$) were isolated from mice bearing SPARC-transduced breast cancer cells (SN25ASP) [1] (authorization number 288/2017-PR) using FACS Aria cell sorter (Becton Dickinson). Anti-tumor N1-neutrophils were isolated from agar plugs as previously described by Sangaletti *et al.* [1].

To achieve transient knock-down of *Zeb1* in murine MDSCs, cells were transfected using a reverse transfection protocol in which siRNAs (Stealth siRNAs MSS210695, MSS210696, MSS210697, Thermo Fisher Scientific) and Lipofectamine 2000 (Thermo Fisher Scientific) have been used. A mix containing 3.25µl of Lipofectamine in 200µl Optimem (Gibco) and another one with 3.25µl of siRNA (20 µM) stock were prepared, and after 5 min at RT were combined and incubated at RT for 30-40 min. 4.5 x $10^5$ MDSCs were seeded and after the incubation, the siRNA/Lipofectamine mix was added. Cells were incubated for 48h and *Zeb1* knock-down and the modulation of selected immunosuppressive genes were assessed using qPCR.

***Human transcriptomic data analysis***

For each experiment raw data were imported in R software, background corrected, log transformed and normalized using Robust Multichip Average (RMA) method from oligo package [2]. Multiple probes representing the same gene were collapsed by selecting the probe with the highest variance across samples through the collapseRows function in the WGCNA package. For DEG analysis, data from GSE6891, GSE12417, GSE15434, GSE16015 and GSE37642, profiled with Human Genome U133 Plus 2.0 Array were selected. Datasets were merged together by matching probes and batch effect was removed through ComBat function from sva package [3]. Samples were separated into two groups according to the median level of ZEB1 expression and DEGs were calculated using the limma package (doi: 10.1093/nar/gkv007) then P values were adjusted for multiple tests using the Benjamini–Hocheberg FDR. Genes with a FDR < 0.05 were considered statistically significant. Pre-ranked GSEA was performed to calculate which hallmark pathways were significantly up or down modulated. Selected genes where charted through a boxplot.

Three datasets including information on survival, GSE6891 (Human Genome U133 Plus 2.0 Array), GSE37642 and GSE12417 (Human Genome U133A Array), were used for molecular and clinical correlation studies. Survival analysis was first performed in each dataset independently and then merging the datasets together as described above. After data quality control, normalization and correction, samples were separated into two groups according to the median level of ZEB1 expression, the Kaplan Meier curves were plotted and the statistical significance was assessed performing a log-rank test.

To assess the differences between diagnosis and relapse we exploited the data from GSE66525, consisting of 11 samples pre and post-chemotherapy. Pre-processed RMA normalized data was downloaded from NCBI Gene Expression Omnibus (GEO) repository and multiple probes representing the same gene were collapsed selecting the probe with the highest variance across samples. Limma package was used to calculate differentially expressed genes and selected genes where charted through a boxplot.

##### *Chromatin Immunoprecipitation (CHIP)*

For Chromatin-Immunoprecipitation, “Simple Chip Chromatin IP Protocol - Magnetic Beads” kit from Cell Signal was used. Briefly, 20*106 cells were cross-linked using 1% formaldehyde at RT for 10 min and quenched in 125 mM glycine for 5 min. According to manufacturer’s instruction, cross-linked chromatin was digested using Micrococcal Nuclease for 20 mins at 37°C and then sonicated to generate DNA fragments averaging 200–500 bp. Samples were incubated O/N at 4 °C with Zeb1 (1 µg, Genetex), or IgG (Cell Signalling) antibodies.

Immunoprecipitated DNA was purified and eluted in 20 µL of H2O and finally analysed by qPCR. Results were expressed as % of Input. For in silico analyses, BAM files were downloaded from ENCODE (https://www.encodeproject.org/) and analysed using Basepair analysis software.

#### Supplementary Figure 1

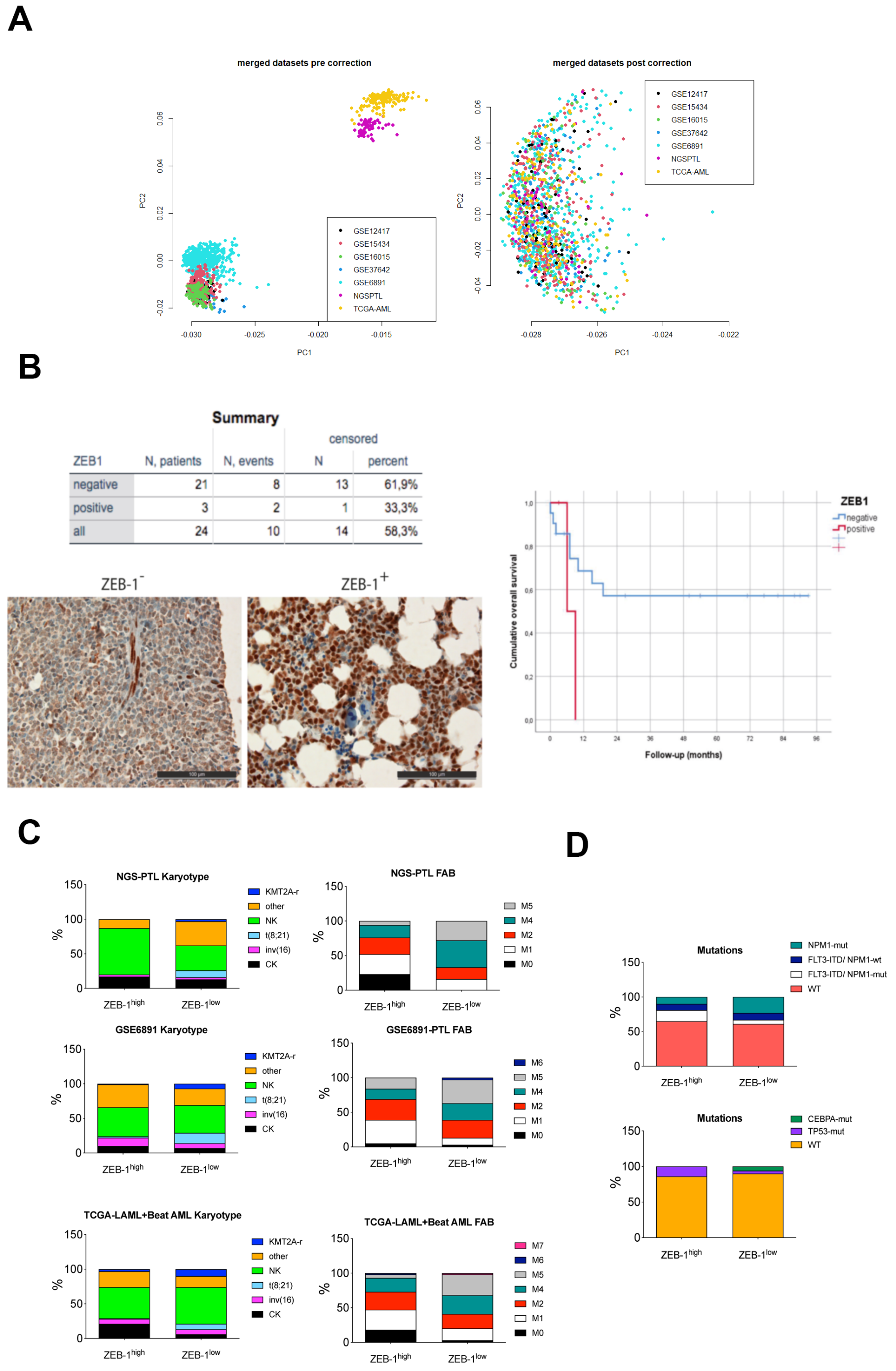

| ZEB1 | N, patients | N, events | censored N | censored percent |
|---|---|---|---|---|
| negative | 21 | 8 | 13 | 61,9% |
| positive | 3 | 2 | 1 | 33,3% |
| all | 24 | 10 | 14 | 58,3% |



**Supplementary Figure 1.** – **A.** OS data from GSE6891, GSE12417, GSE15434, GSE16015, NGS-PTL, TCGA and GSE37642 were merged together by matching probes and batch effect was removed through ComBat function from sva package. B. ZEB-1 expression levels in a cohort of

24 AML patients (ZEB1$^{high}$ n=3 and ZEB1$^{low}$ n=21) detected by IHC analysis and Kaplan-Meier curves showing the reduced overal survival of ZEB-1$^{high}$ (positive) patients compared with ZEB-1$^{low}$ (negative) patients. C. Cytogenetic characteristics and FAB of AML patients included in our cohort (NGS-PTL, karyotype, ZEB1$^{high}$ n=30 and ZEB1$^{low}$ n=31; FAB, ZEB1$^{high}$ n=17 and ZEB1$^{low}$ n=18), GSE6891 (karyotype, ZEB1$^{high}$ n=206 and ZEB1$^{low}$ n=208; FAB, ZEB1$^{high}$ n=219 and ZEB1$^{low}$ n=233) and TCGA-LAML+Beat AML (karyotype, ZEB1$^{high}$ n=138 and ZEB1$^{low}$ n=140; FAB, ZEB1$^{high}$ n=85 and ZEB1$^{low}$ n=103) public datasets between ZEB1$^{high}$ and ZEB1$^{low}$ AML cases. ZEB1$^{high}$ and ZEB1$^{low}$ AML patients were dichotomized according to the median expression of ZEB-1. **D.** Percentage of patients ZEB1$^{high}$ and ZEB1$^{low}$ AML cases from the TCGA-LAML+Beat AML datasets (ZEB1$^{high}$ n=133 and ZEB1$^{low}$ n=137) carrying the selected mutations.

#### Supplementary Figure 2

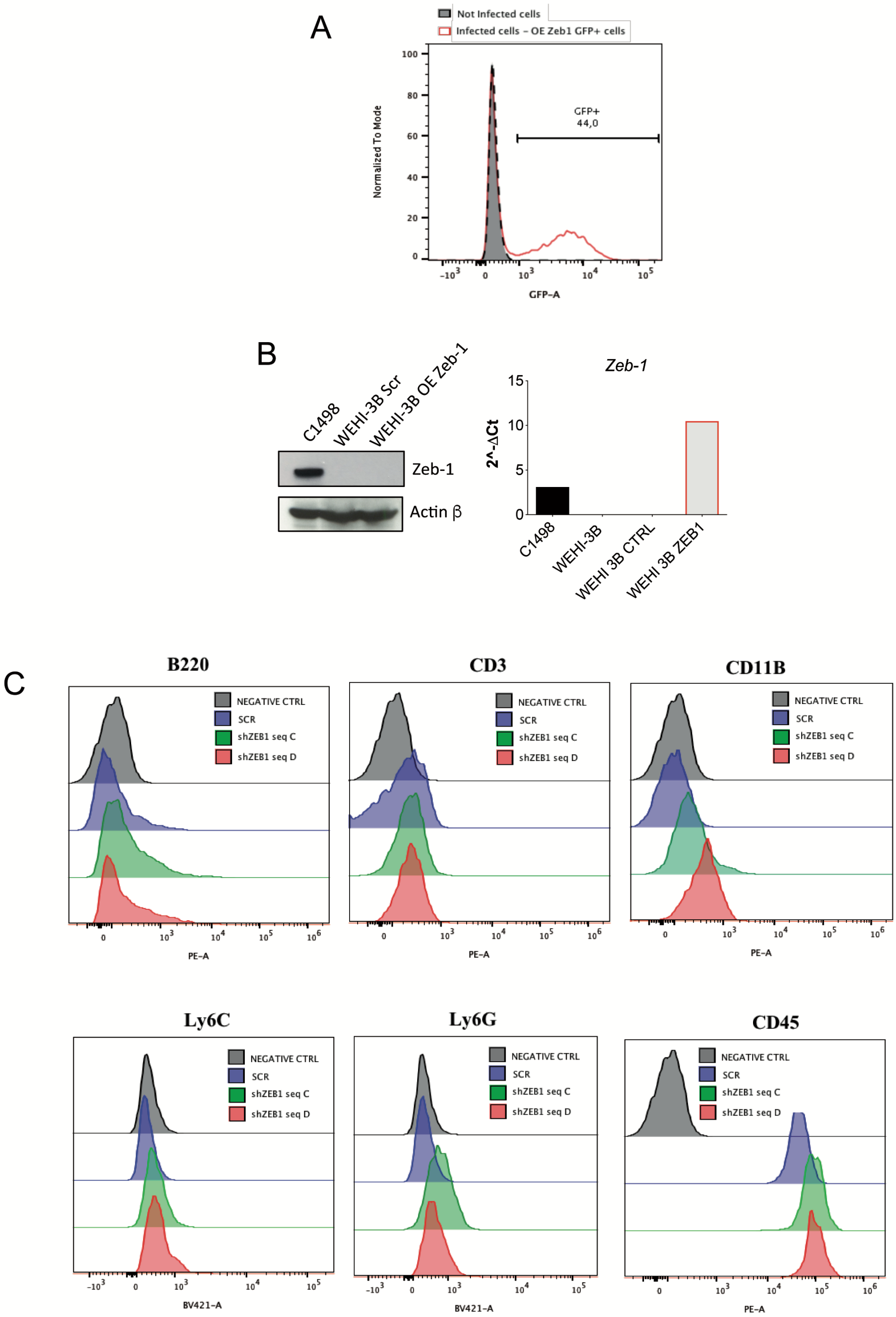

**Supplementary Figure 2.** –**A.** FACS analysis showing the percentage of WEHI-3B GFP+ cells after infection to overexpress (OE) Zeb1. **B.** Western blot and qPCR showing Zeb1 levels in WEHI-3B OE Zeb1 and parental cell line. C1498 cells were used as positive controls. **C.** Flow cytometry analysis showing the expression of B220, CD3, CD11b, Ly6C, Ly6G and CD45 in term of MFI in C1498 cells infected with shZeb1 seq C and seq D and Scramble cells. Unstained cells were used as negative control.

#### Supplementary Figure 3

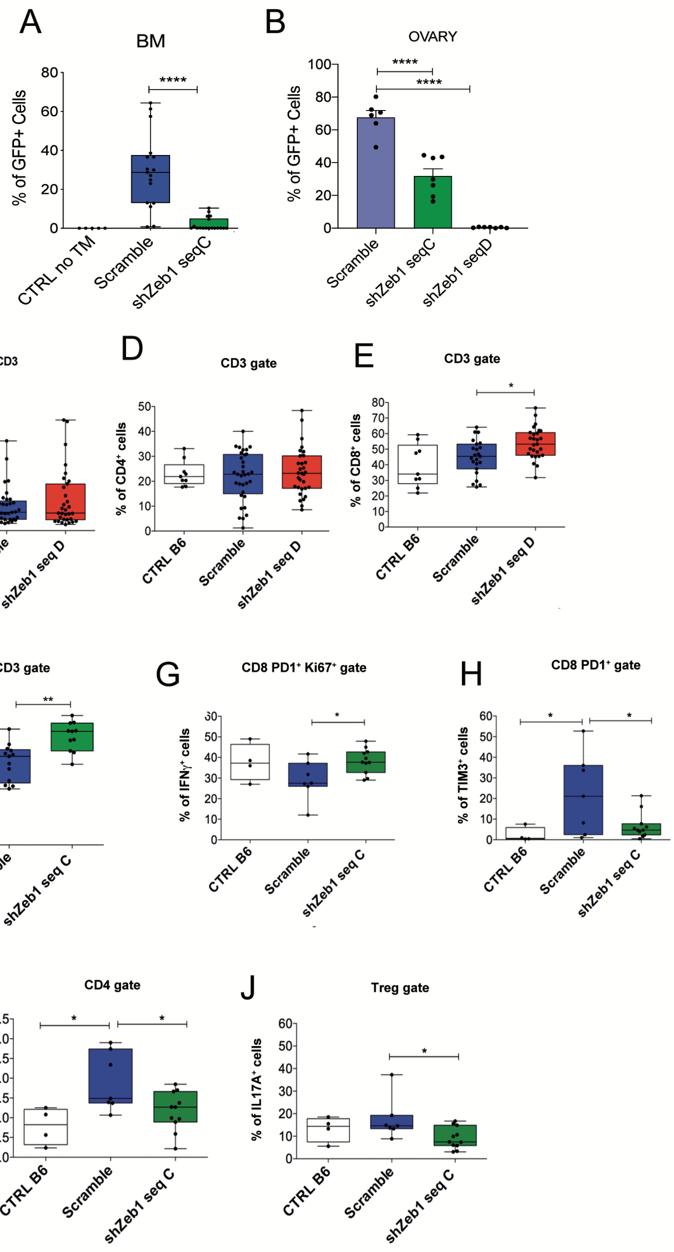


**Supplementary Figure 3.** – **A.** Frequency of GFP+ cells within the BM of mice injected i.b. with Scramble (n=17) and shZeb-1 seq C (n=18) cells. Data are representative of a pool of 2 independent experiments. Mice were sacrificed after 34 days and naïve C57BL/6mice were used as control (B6 CTRL n=5). Statistical analysis: One way ANOVA; the p value relative to the comparison Scramble vs shZeb1 seq C is shown in the figure ****p<0.0001; KW test p<0.0001 **B.** Frequency of GFP+ cells within the ovary of mice injected with Scramble (n=6), shZeb1 seq C (n= 7), or shZeb1 seq D (n= 7). Data are representative of an individual experiment out of 2 performed (Statistical analysis: One way ANOVA followed by Tukey's multiple comparison test, p<0.0001; the p value relative to the comparison Scramble vs shZeb1 seq C or seq D is shown in the figure (****p<0.0001) **C.** Frequency of CD3+ cells within the BM of mice injected i.b. with Scramble (n=33) and shZeb-1 seq D (n=31) cells. Naïve C57BL/6mice were used as control (B6 CTRL n=9). Data represent a pool of 3 independent experiments **D.** Frequency of CD4+ cells within the BM of mice injected i.b. with Scramble (n=33) and shZeb-1 seq D (n=31) cells. Naïve C57BL/6mice were used as control (B6 CTRL n=9). Data represent a pool of 3 independent experiments **E.** Frequency of CD8+ cells within the BM of mice injected i.b. with Scramble (n=23) and shZeb-1 seq D (n=27) cells. Naïve C57BL/6mice were used as control (B6 CTRL n=9). Data represent a pool of 2 independent experiments. Statistical analysis: One-way ANOVA followed by Tukey's multiple comparison test p=0.0014; the p value relative to the comparison Scramble vs shZeb1 seq D is shown in the Figure (*p<0.05) **F.** Frequency of CD8+ cells within the BM of mice injected i.b. with Scramble (n=13) and shZeb-1 seq C (n=11) cells. Naïve C57BL/6mice were used as control (B6 CTRL n=9). Data are representative of an individual experiment out of 2 performed; Statistical analysis: One way ANOVA followed by Tukey's multiple comparison test, p<0.0001; the p value relative to the comparison Scramble vs shZeb1 seq C is shown in the figure (**p<0.01) **G.** Frequency of activated CD8+PD1+Ki67+IFN-γ cells and **H.** exhausted T-cells (CD8+TIM3+PD1+) within the BM of mice injected i.b. with Scramble (n=7) and shZeb-1 seq C (n=11) cells. Mice were sacrificed after 34 days and naïve C57BL/6mice were used as control (B6 CTRL n=4). Data are representative of an individual experiment out of 2 performed. Statistical analysis: One way ANOVA followed by Tukey's multiple comparison test; p=0.0212; the p value relative to the comparison Scramble vs CTRL B6 or shZeb1 seq C is shown in the figure (*p<0.05) **I.** Frequency of IL-17A producing CD4+ (Statistical analysis: One-way ANOVA followed by Tukey's multiple comparison test; p=0.0122; the p value relative to the comparison Scramble vs CTRL B6 or shZeb1 seq C is shown in the figure (*p<0.05)) and **J.** Treg (CD4+CD25+Foxp3+) cells within the BM of mice injected i.b. with Scramble (n=7) and shZeb-1 seq C (n=11) cells. Mice were sacrificed after 34 days and naïve C57BL/6mice were used as control (B6 CTRL n=4). Data are representative of an individual experiment out of 2 performed. (Statistical analysis: One-way ANOVA followed by Tukey's multiple comparison test; p=0.0576; the p value relative to the comparison Scramble vs shZeb1 seq C is shown in the figure (*p<0.05))

#### Supplementary Figure 4

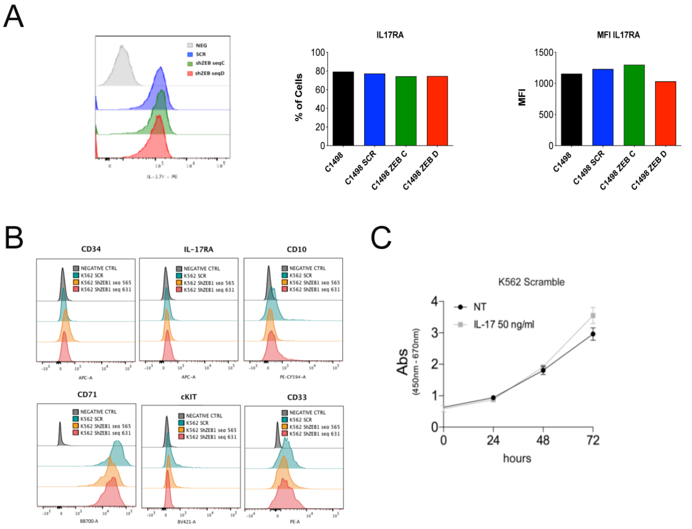


**ENCFF070VUK** TF ChIP-seq GM12878 *ZEB1*

D

| Chromosome | Start | End | Lenght | Tags | -10 Log (pval) | FDR (%) | Fold | Annotation | Accession | Symbol |
|---|---|---|---|---|---|---|---|---|---|---|
| chr12 | 93,569,512 | 93,571,589 | 2,078 | 146 | 317.01 | NaN | 14.96 | Promoter | ENST00000548537.1;ENST00000551556.1;ENST00000499137.6;ENST00000549122.5;ENST00000547845.5;ENST00000551626.1;ENST00000549887.1;ENST00000340600.6;ENST00000548091.5;ENST00000536696.6;ENST00000500986.5;ENST00000622746.4;ENST00000549206.5;ENST00000547229.1 | **SOCS2-AS1;SOCS2** |
| chr19 | 41,341,470 | 41,342,080 | 611 | 28 | 132.76 | NaN | 18.17 | Promoter | ENST00000598758.5 | **TGFB1** |
| chr19 | 41,352,296 | 41,353,352 | 1,057 | 41 | 58.45 | NaN | 7.74 | Promoter | ENST00000604123.5;ENST00000539627.5;ENST00000604424.1;ENST00000221930.5 | **TGFB1;AC011462.1;TMEM91** |
| chr19 | 41,353,434 | 41,354,416 | 983 | 46 | 89.27 | NaN | 6.45 | Promoter | ENST00000604123.5;ENST00000221930.5 | **TGFB1;TMEM91** |
| chr12 | 56,338,615 | 56,339,128 | 514 | 43 | 202.29 | NaN | 19.69 | Promoter | ENST00000228534.5 | **IL23A** |

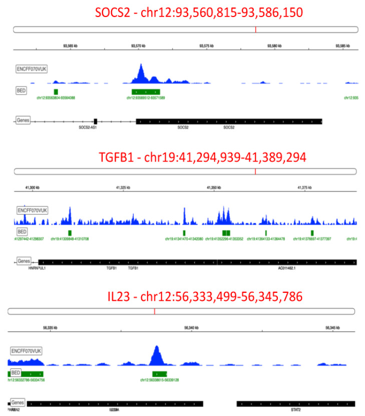


**Supplementary Figure 4.** – **A.** FACS analysis showing IL-17RA levels in term of MFI and percentage of positive cells among the Zeb-1 expressing or silenced C1498 cells. **B.** Flow cytometry analysis showing the expression of CD34, IL-17RA, CD10, CD71, cKIT and CD33 in term of MFI in C1498 cells infected with shZEB1 seq C and seq D and Scramble cells. Unstained cells were used as negative control. **C.** Cell proliferation assay performed on ZEB-1 expressing (Scramble) cells upon IL-17A stimulation (50 ng/mL) assessed by Xtt assay after 24h, 48h and 72h. Cell proliferation for each time point was calculated as the (Absorbance (Abs) at 450 nm – Abs at 670 nm). **D.** ChIP-seq profile of ZEB1 on *TGFB1*, *SOCS2* and *IL23A* loci in GM12878 cells. **F.** Heatmap of canonical pathway enrichment analysis showing different immunological features between *ZEB1*$^{high}$ and *ZEB1*$^{low}$

**Supplementary Table 1**

| Gene Symbol | Assay ID | species reactivity |
|---|---|---|
| Zeb1 | Mm00495564_m1 | mouse |
| Arg1 | Mm00475988_m1 | mouse |
| Il23a | Mm00518984_m1 | mouse |
| Tgfb1 | Mm01178820_m1 | mouse |
| Il6 | Mm00446190_m1 | mouse |
| Il2 | Mm00434256_m1 | mouse |
| Socs2 | Mm00850544_g1 | mouse |
| Mmp9 | Mm00442991_m1 | mouse |
| Actb | Mm02619580_g1 | mouse |
| ZEB1 | Hs00232783_m1 | human |
| TGFB1 | Hs00998133_m1 | human |
| IL23A | Hs00372324_m1 | human |
| IL6 | Hs00174131_m1 | human |
| SOCS2 | Hs00919620_m1 | human |
| MMP9 | Hs00957562_m1 | human |
| ACTB | Hs03023943_g1 | human |

| antibodies | clone | fluorochrome | species reactivity | company | application |
|---|---|---|---|---|---|
| CD11b | M1/70 | BB700 | mouse | BD | FC |
| Ly6G | 14S-2C11 | BV421 | mouse | BD | FC |
| Ly6C | AL-21 | BV605 | mouse | BD | FC |
| CD3 | 145-2C11 | BV786 | mouse | BD | FC |
| CD4 | GK1.5 | PE | mouse | eBioscience | FC |
| OX40 | OX-86 | APC | mouse | eBioscience | FC |
| CD8 | 53-6.7 | APC-Cy7 | mouse | BioLegend | FC |
| CD25 | PC61 | BV650 | mouse | BD | FC |
| Foxp3 | FJK-16S | PerCp-Cy5.5 | mouse | eBioscience | FC |
| Ki-67 | B56 | Alexa 647 | mouse | BD | FC |
| PD1 | RMP1-30 | BV421 | mouse | BD | FC |
| IFN-γ | XMG1.3 | PE-CF595 | mouse | BD | FC |
| TNFα | MP6-XT22 | BV510 | mouse | BD | FC |
| IL17A | TC11-18H10 | FITC | mouse | BioLegend | FC |
| TIM3 | SD12/TIM3 | BV421 | mouse | BD | FC |
| ZEB1 (E2G6Y) | | | human, mouse, rat | Cell Signaling Technology | WB |
| β-Actin | | | human, mouse, rat | Sigma | WB |

**Supplementary Table 1 -** List of Taqman probes used in qPCR experiments for murine and human cell lines and antibodies used for flow cytometry for in vivo experiments and western blot experiments

### Supplementary Table 2

| cohort | ID | Gender | Disease | Disease.type | Disease_Stage | specimen_Type | Age_at_Dx | Karyotype | cytogenetic.risk | trisomy | FAB | WBC_count | TypeTreatment |
|---|---|---|---|---|---|---|---|---|---|---|---|---|---|
| **NGS-PTL** | 1 | FEMALE | AML | de_novo | dx | BM | 54 | other | intermediate | n | NA | 38,9 | intensive |
| **NGS-PTL** | 3 | MALE | AML | de_novo | dx | BM | 66 | other | intermediate | n | M4 | 115 | intensive |
| **NGS-PTL** | 5 | FEMALE | AML | de_novo | dx | BM | 68 | other | adverse | n | NA | 8,6 | not_treated |
| **NGS-PTL** | 6 | MALE | AML | de_novo | dx | BM | 57 | normal | intermediate | n | M4 | 2,9 | not_treated |
| **NGS-PTL** | 7 | FEMALE | AML | de_novo | dx | BM | 59 | normal | intermediate | n | M2 | 1,6 | intensive |
| **NGS-PTL** | 8 | FEMALE | AML | NA | dx | BM | NA | normal | intermediate | n | NA | NA | NA |
| **NGS-PTL** | 9 | FEMALE | AML | t-AML | dx | BM | 74 | complex | adverse | y | M5 | 2,8 | intensive |
| **NGS-PTL** | 11 | MALE | AML | de_novo | dx | BM | 65 | other | intermediate | n | NA | 46,5 | intensive |
| **NGS-PTL** | 12 | FEMALE | AML | t-AML | dx | BM | 76 | other | intermediate | n | NA | NA | NA |
| **NGS-PTL** | 13 | FEMALE | AML | NA | dx | BM | 69 | complex | adverse | n | M4 | NA | NA |
| **NGS-PTL** | 14 | FEMALE | AML | de_novo | dx | BM | 51 | normal | intermediate | n | M0 | 3,8 | intensive |
| **NGS-PTL** | 15 | FEMALE | AML | de_novo | dx | BM | 47 | normal | intermediate | n | M2 | 89,4 | intensive |
| **NGS-PTL** | 16 | MALE | AML | de_novo | dx | BM | 67 | normal | intermediate | n | M4 | 9,7 | intensive |
| **NGS-PTL** | 18 | FEMALE | AML | NA | dx | BM | 42 | normal | intermediate | n | NA | 14,5 | NA |
| **NGS-PTL** | 20 | MALE | AML | sec | dx | BM | 71 | normal | intermediate | n | NA | 2,3 | intensive |
| **NGS-PTL** | 21 | FEMALE | AML | de_novo | dx | BM | 70 | other | intermediate | y | M4 | NA | intensive |
| **NGS-PTL** | 23 | MALE | AML | de_novo | dx | BM | 82 | other | intermediate | n | NA | NA | not_intensive |
| **NGS-PTL** | 24 | MALE | AML | t-AML | dx | BM | 69 | complex | adverse | n | M5 | 238 | not_treated |
| **NGS-PTL** | 25 | MALE | AML | de_novo | dx | BM | 62 | complex | adverse | n | M0 | 1,5 | intensive |
| **NGS-PTL** | 26 | FEMALE | AML | de_novo | dx | BM | 67 | normal | intermediate | n | M0 | 108,6 | intensive |
| **NGS-PTL** | 34 | MALE | AML | NA | dx | BM | NA | other | adverse | n | NA | NA | NA |
| **NGS-PTL** | 35 | MALE | AML | t-AML | dx | BM | 62 | complex | adverse | y | NA | 2,7 | NA |
| **NGS-PTL** | 36 | MALE | AML | NA | dx | BM | NA | complex | adverse | NA | NA | NA | NA |
| **NGS-PTL** | 37 | MALE | AML | de_novo | dx | BM | 39 | t(8;21) | favorable | n | M2 | NA | intensive |
| **NGS-PTL** | 38 | FEMALE | AML | de_novo | dx | BM | 62 | MLL-rearranged | intermediate | y | M5 | 23,1 | intensive |
| **NGS-PTL** | 39 | FEMALE | AML | de_novo | dx | BM | 50 | normal | intermediate | n | M5 | 77,7 | intensive |
| **NGS-PTL** | 40 | FEMALE | AML | NA | dx | BM | 76 | normal | intermediate | n | NA | NA | NA |
| **NGS-PTL** | 41 | FEMALE | AML | de_novo | dx | BM | 60 | normal | intermediate | n | NA | 68,5 | not_treated |
| **NGS-PTL** | 43 | FEMALE | AML | t-AML | dx | BM | 62 | other | intermediate | n | M1 | 13,4 | intensive |
| **NGS-PTL** | 44 | MALE | AML | de_novo | dx | BM | 66 | normal | intermediate | n | M4 | 18,9 | intensive |
| **NGS-PTL** | 45 | MALE | AML | de_novo | dx | BM | 42 | normal | intermediate | n | NA | 163,9 | not_treated |
| **NGS-PTL** | 46 | MALE | AML | de_novo | dx | BM | 45 | normal | intermediate | n | M5 | 88 | intensive |
| **NGS-PTL** | 47 | FEMALE | AML | de_novo | dx | BM | 66 | normal | intermediate | n | NA | 35,9 | intensive |
| **NGS-PTL** | 48 | FEMALE | AML | de_novo | dx | BM | 60 | normal | intermediate | n | M1 | 3,2 | intensive |
| **NGS-PTL** | 49 | FEMALE | AML | de_novo | dx | BM | 72 | normal | intermediate | n | NA | 26,1 | not_treated |
| **NGS-PTL** | 50 | FEMALE | AML | de_novo | dx | BM | 34 | normal | intermediate | n | M1 | 102 | intensive |
| **NGS-PTL** | 51 | FEMALE | AML | de_novo | dx | BM | 38 | normal | intermediate | n | M1 | 37,2 | intensive |
| **NGS-PTL** | 53 | FEMALE | AML | NA | dx | BM | NA | complex | adverse | NA | M2 | NA | NA |
| **NGS-PTL** | 54 | MALE | AML | de_novo | dx | BM | 61 | inv(16)/t(16;16) | favorable | n | M4 | 7,4 | intensive |
| **NGS-PTL** | 55 | FEMALE | AML | de_novo | dx | BM | 71 | other | intermediate | y | M4 | 90 | intensive |
| **NGS-PTL** | 56 | FEMALE | AML | t-AML | dx | BM | 62 | complex | adverse | n | NA | 77 | not_treated |
| **NGS-PTL** | 57 | MALE | AML | t-AML | dx | BM | 68 | other | adverse | n | M0 | 5,2 | not_intensive |
| **NGS-PTL** | 58 | MALE | AML | de_novo | dx | BM | 42 | other | intermediate | y | M5 | 66,9 | intensive |
| **NGS-PTL** | 59 | MALE | AML | de_novo | dx | BM | 64 | normal | intermediate | n | M1 | 1,9 | intensive |
| **NGS-PTL** | 60 | MALE | AML | de_novo | dx | BM | 64 | normal | intermediate | n | M1 | 189,5 | NA |
| **NGS-PTL** | 61 | MALE | AML | de_novo | dx | BM | 77 | normal | intermediate | n | NA | 6,7 | not_intensive |
| **NGS-PTL** | 64 | FEMALE | AML | sec | dx | BM | 66 | normal | intermediate | n | NA | NA | NA |
| **NGS-PTL** | 65 | MALE | AML | de_novo | dx | BM | 64 | normal | intermediate | n | M1 | 65,1 | intensive |
| **NGS-PTL** | 66 | MALE | AML | de_novo | dx | BM | 70 | normal | intermediate | n | NA | 234 | intensive |
| **NGS-PTL** | 67 | MALE | AML | NA | dx | BM | NA | normal | intermediate | n | NA | NA | NA |
| **NGS-PTL** | 68 | FEMALE | AML | t-AML | dx | BM | 57 | inv(16)/t(16;16) | favorable | n | M4 | 10,5 | intensive |
| **NGS-PTL** | 69 | FEMALE | AML | de_novo | dx | BM | 72 | other | intermediate | n | M2 | 35,1 | not_treated |
| **NGS-PTL** | 70 | MALE | AML | de_novo | dx | BM | 31 | t(8;21) | favorable | n | M4 | 5,1 | intensive |
| **NGS-PTL** | 71 | MALE | AML | de_novo | dx | BM | 67 | complex | adverse | n | M2 | 3,6 | intensive |
| **NGS-PTL** | 72 | MALE | AML | de_novo | dx | BM | 52 | t(8;21) | favorable | n | NA | NA | intensive |
| **NGS-PTL** | 74 | MALE | AML | sec | dx | BM | 66 | other | intermediate | n | NA | 9,6 | not_intensive |
| **NGS-PTL** | 75 | FEMALE | AML | de_novo | dx | BM | 39 | other | intermediate | n | M1 | 43,2 | intensive |
| **NGS-PTL** | 83 | MALE | AML | de_novo | dx | BM | 73 | normal | intermediate | n | NA | 2 | not_intensive |
| **NGS-PTL** | 85 | FEMALE | AML | de_novo | dx | BM | 67 | normal | intermediate | n | NA | 46,7 | intensive |
| **NGS-PTL** | 100 | MALE | AML | sec | dx | BM | 66 | normal | intermediate | n | NA | 3,9 | not_intensive |
| **NGS-PTL** | 101 | FEMALE | AML | de_novo | dx | BM | 63 | normal | intermediate | n | M2 | 44,3 | intensive |

**NA:** not available

**Supplementary Table 2** – Clinical features of AML patients included in NGS-PTL cohort

| Dataset ID | Platform | Sample n. and type | Normalization | Data analysis |
|---|---|---|---|---|
| GSE6891, GSE12417, GSE15434, GSE16015 | Human Genome U133 Plus 2.0 | 973 AML BM, PB MNCs | sst-RMA | DEG between $ZEB1^{high}$ and $ZEB1^{low}$ |
| NGS-PTL (GSE161532) | Affymetrix HTA 2.0 | 61 AML BM MNCs (blasts ≥80%) | sst-RMA | karyotype, FAB, OS |
| GSE6891 | Affymetrix U133 Plus 2.0 | 488 AML MNCs | RMA | karyotype, FAB, OS |
| GSE37642 | Affymetrix U133A | 405 AML BM MNCs | RMA | FAB, OS |
| TCGA-LAML + Beat AML | RNA-seq | 284 BM MNCs | CPM | karyotype, FAB, OS, mutations |
| GSE66525 | Affymetrix ST1.1 | Paired PB samples of 11 AML patients at Dx and Rel | RMA | |

BM: bone marrow, CPM: Counts Per Million; DEG: differentially expressed genes; FAB: French-American-British classification; MNCs: mononuclear cells; OS: overall survival; RMA: Robust Multi-Array average; sst-RMA: signal space transformation robust multi-array average; TMM: Trimmed Mean of M values.

**Supplementary Table 3** – Human dataset included in the manuscript
